# The probable numbers of kin in a multi-state population: a branching process approach

**DOI:** 10.64898/2026.03.31.715515

**Authors:** Joe W. B. Butterick

## Abstract

Recent progress in mathematical kinship modelling has allowed one to predict the probable numbers of kin for a typical population member. In the models, kin may be structured by age and sex, both in static or time-variant demographies. Knowing the probable numbers of kin in different *stages* – such as parity, health status, or geographic location – however, remains an open challenge in Kinship Demography. Knowing how population structure delimits kin to distinct stages is an advance – for instance, the probability of having one sister at home and one sister away has different social implications from the probability of having two sisters.

We present a novel analytical framework, grounded in branching process theory, that provides kin-number distributions jointly structured by age and stage. Using recursive compositions of probability generating functions (PGFs), we derive the joint age, stage, and age × stage kin-number distributions. All marginal distributions over either dimension naturally emerge. Simple extensions of the PGF approach additionally yield: the joint distribution of an individual’s own stage and their kin’s stage; the probable numbers of kin deaths, both in total and by generation number; and the probabilities of being kinless and/or orphaned. We demonstrate the framework through novel results in an application using UK parity-specific fertility and mortality data.

**Highlights:**

- A new method calculates probability generating functions for the number of kin structured by age and stage
- The model allows predicting the probable numbers of kin organised by age and stage
- Recursive nesting of probability generating functions in branching processes is used
- An application is presented highlighting the novel results

## 1. Introduction

Kinship demography is an expanding area of research. Applications of the topic span fields ranging from population ecology [1, 2, 3] to human demography [4, 5]. The number of kin an individual has is determined by its reproduction and the reproduction of others in its so called kin-network. Within the context of ecological systems, and in particular inclusive fitness theory [6], the size of an animal’s related kin-network has been shown to influence its propensity to demonstrate cooperative behaviour across its life-course [7, 8]. In humans, kin availability has consequences for care systems [9], emotional and financial support [10], mental state [11, 12], and social and economic inequalities between and within generations [13, 14].

The mathematical foundations for predicting the availability of ones kin were set by the seminal work of Goodman, Keyfitz & Pullum [15]. The authors therein proposed a method to calculate the expected numbers of ancestors and siblings. More recently, there has been a comprehensive expansion in mathematical models, in large part motivated by the pioneering framework of Caswell [16]. The author uses a system of matrix projections to predict, for a typical population member called “Focal” (in this manuscript also we refer to this population member as Focal), the expected numbers of kin structured by their ages. Extensive developments of the model, led by Caswell and co-workers, have allowed matrix projections to provide the expected numbers of kin structured by sex [17], stage [18], and in time-variant demographies [19]. Such progress has, for example, allowed one to predict the future kin numbers across all countries in the world [20]. Concomitant to the advances made by Caswell and collaborators, well-considered mathematical models proposed by Coste *et al*. [1] and Coste [21] have provided the expected numbers of kin under any population structure. With a broader emphasis – also focusing on population ecology – the research of Coste and co-authors complements the work of Caswell through providing the benefits of being able to predict the numbers of kin structured by class, spatial distance, or dispersal.

Notwithstanding such remarkable advances in kinship modelling, each above-mentioned model predicts the expected numbers of kin (but see Caswell [22] who also extracts the variance). In reality kin-numbers are non-negative integers and are better represented through probability mass functions (PMFs). Butterick *et al*. [23] provide the first probabilistic representation of the kin-network in a one-sex time-invariant demography. Inspired by the characterisation of the branching-process of kinship [24, 25], the authors show, using three concise formulae, how to calculate for each age of a typical population member, the probabilities that they have *j* = 0, 1, …, of a certain kin-type of given age (or range of ages). Building on this approach, Butterick [26] extends the model to calculate the probable numbers of kin by age, sex, and time. An extension of the latter model also calculates the probabilities of kin-loss over the life course.

Having a distribution for kin-number offers several advantages over knowing only the expected numbers. For instance, the full distribution allows for calculation of higher order moments, such as the skew and kurtosis, important measures in the study of lifetime reproductive output [27, 28]. Additionally, a complete number distribution allows one to condition on the probabilities of having so many kin with specific needs [29, 9]. Extending these probabilistic formulations to stage-structured populations is, however, non-trivial. In this research, we propose a novel method to do so. By drawing from the theory of branching processes, but applying to probability generating functions rather than probability mass functions, here we present a framework to derive the probable numbers of kin structured by stage and age.

Branching processes are naturally appealing to model kinship; they offer the ability to recursively map generation-on-generation replenishment. For example, suppose the random variable *X* represents the number of offspring an individual has over their life and suppose the random variable *Y*_*t*_ represents population size at time *t*. Let 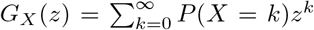 the probability generating function (PGF) for *X*, and let 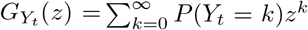 the generating function for *Y*_*t*_. The recursion 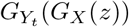 generates the population size of the next generation *t* + 1. Generating functions were used to estimate kin numbers in a variety of mathematical papers published in the 1980s. By composing PGFs in a Galton-Watson process, Waugh [25] and Joffe & Waugh [30] extracted kin-numbers unstructured by age (in non-overlapping generations), while Joffe & Waugh [31] extended the method using multi-type Galton-Watson processes with age in lieu of type (in overlapping generations). In continuous time, and again accounting for overlapping generations, so-called Crump-Mode-Jagers processes (named after Crump & Mode [32] and Jagers [33]) paved way for solving problems such as birth rank and sib-ship size [24, 34], although in stable populations and not in readily generalisable frameworks. This research is motivated by the branching process method. In what follows, we recursively compose PGFs to calculate the probable numbers of kin in a time-invariant, one-sex population, structured by age *and* stage. We demonstrate its application using an example of parity in the UK.

## 2. Model

This model extends, to a multi-state demography, recent research [23, 26] which derive kin-number distributions in an age-structured populations. Here, we work directly with PGFs rather than the probability mass functions. In Appendix B we show, when removing stage, the present model is equivalent to the above-mentioned frameworks.

### 2.1. Demography and assumptions

We consider a population structured by age-classes *a* = 0, 1, …, *ω* and stages *s* = 1, …, *k*. Let 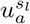 represent the probability that an individual survives, starting from stage *s*_*l*_, from age-class *a* to age-class *a* + 1. Let 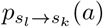 be the probability that between age-classes *a* and *a* + 1 the individual moves from stage *s*_*l*_ to *s*_*k*_. Let 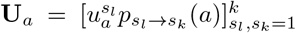 be a matrix with entries the age-class-specific survival and stage-transition probabilities. Let 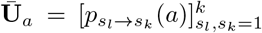 be a (row stochastic) matrix with entries the age-specific stage-transitions. Let 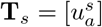 be a matrix with entries the stage-specific survival probabilities.

We assume independence in reproduction across ages-classes. That is, the number of offspring born to an individual in age-class *a*′ is independent of the number born in age-class *a*^*′′*^. Let 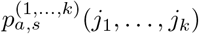 be the probability that an individual of age-class *a* and stage *s* produces *j*_1_, …, *j*_*k*_ offspring into stages 1, …, *k*. For each age-class *a*, we use this to derive a matrix of expected rates at which individuals of stage *s*_*a*_ produce offspring of stage *s*_0_. Let **F**_*a*_ have (*i, l*) entries the expected rate at which stage *i* age *a* produce stage *l* (i.e., 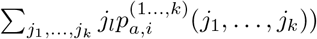. Let **D**_*s*_ distribute newborns of stage *s* into appropriate age-classes (in this framework, the first age-class).

Following the approach of Caswell [18], we construct a multi-state projection matrix **Ã**. Let 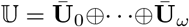 and 𝔽 = **F**_0_ ⊕ · · · ⊕ **F**_*ω*_ account for survival and fertility. Let 𝕋 = **T**_1_ ⊕ · · · ⊕ **T**_*k*_ and ℍ= **H**_1_ ⊕ · · · ⊕ **H**_*k*_ account for stage-transitions and the redistribution of newborns. Using the commutation matrix **Ψ**_*q,r*_ (defined through **Ψ**_*q,r*_vec**A** = vec**A**^*†*^ for **A** ∈ *M*_*q,r*_ [35]), define

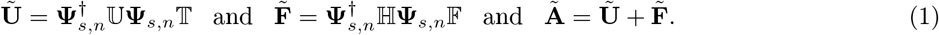

(see Caswell [18] for further details). The block-structured projection matrix, as will be seen in Section 3.2.1, is required to infer the paternal state distributions of Focal’s ancestors. Here, state within the multi-state framework is defined through the vec-permutation approach, encompassing both age *a* and stage *s* through the relation *ka* + *s*. Moving forwards, sometimes we will denote state by (*a, s*).

**Table 1:**
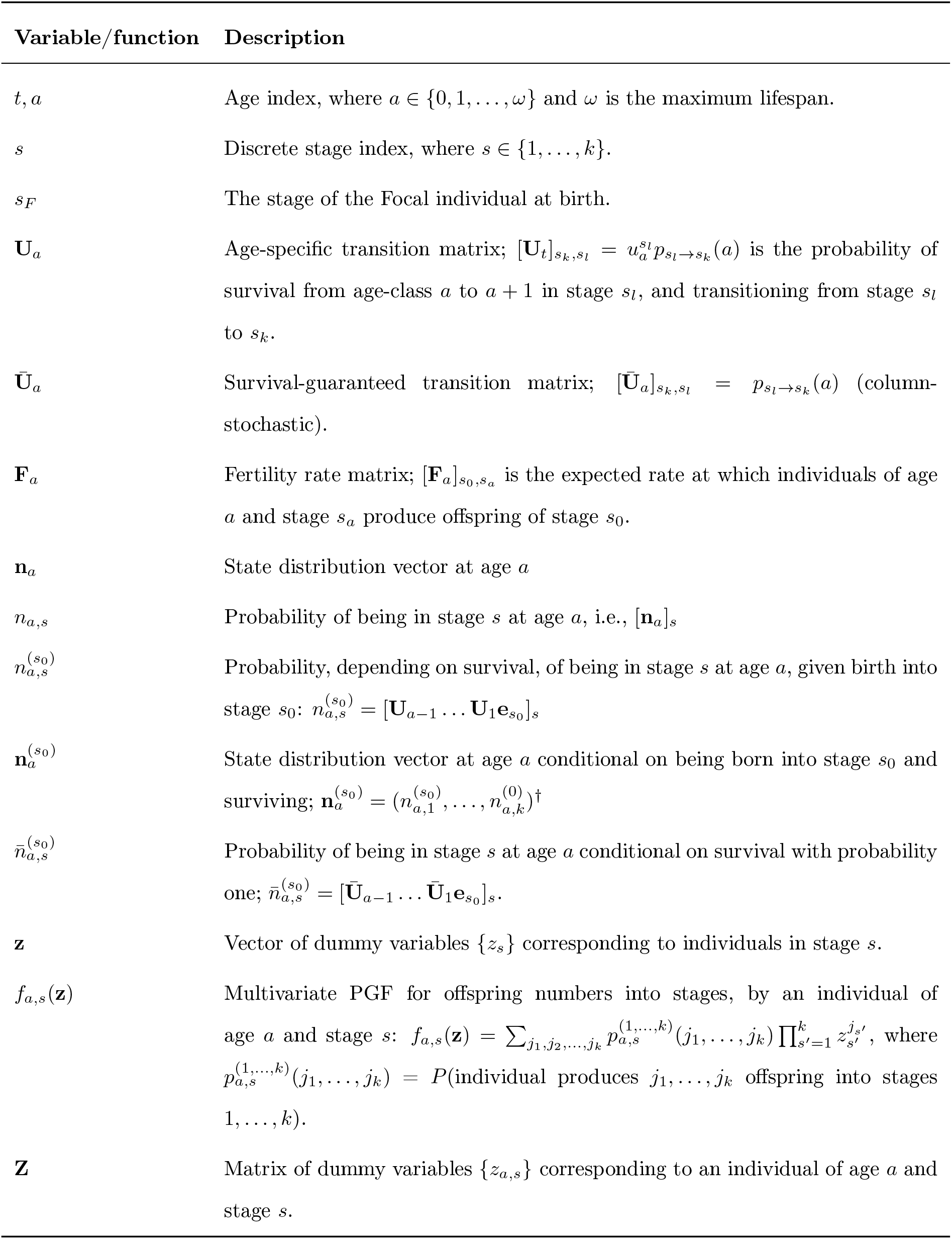
Variables and assumptions.

### 2.2. Kin state distribution PGF

For an individual born in stage *s*_0_, we define their *state distribution* PGF to describe their state *t* time-steps after birth:

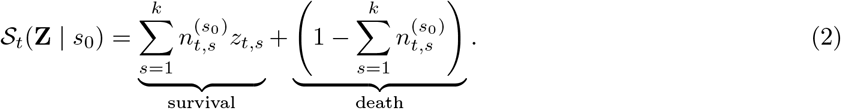

In Eq (2), the term under-braced “survival” tracks the probability the individual is alive and in stage *s* at age *t*, counted using the dummy variable *z*_*t,s*_, while the term under-braced “death” accounts for the probability that the individual has died by age *t* (effectively normalising the PGF). As a simple interpretation of how Eq (2) operates, consider a newborn cohort. Let 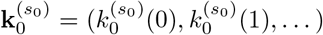 be a (column) vector representing the probable numbers *j* = 0, 1, …, of newborns born into state *s*_0_. In PGF form we write 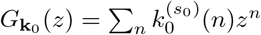. Suppose we *only* wish to know the probable numbers of the cohort who reach age *t*, and are by that time in stage *s*. By setting 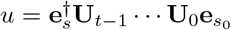 we reduce Eq (2) to 𝒮_*t*_(*z*_*t,s*_ | *s*_0_) = (1 − *u*) + *uz*_*t,s*_, which is the PGF for the Bernoulli random variable *u*. The PGF encodes the probability an individual survives to stage *s* at time *t*. By substituting 𝒮_*t*_(*z*_*t,s*_ | *s*_0_) for *z* in 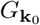 we obtain

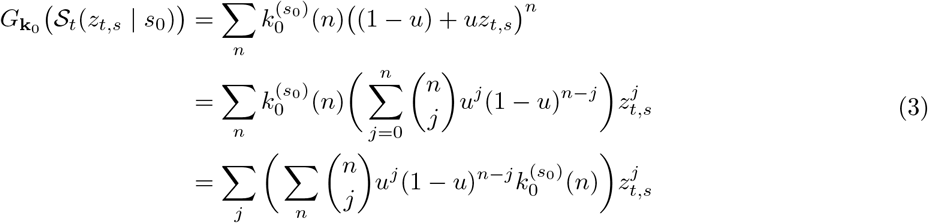

from which we find the probable numbers of the newborn cohort who reach stage *s* by age *t*, by extracting the *s, t* coefficient of the Taylor expansion of 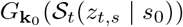:

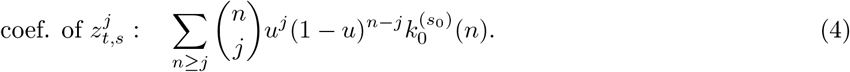

We interpret Eq (4) as follows: out of *n* Bernoulli trials, there are *j* successes (and *n* − *j* failures) – each trial represents the life trajectory of an independent individual; success represents the trajectory ending in stage *s*. By substituting the PGF 𝒮_*t*_(**Z** | *s*_0_) into 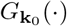, the multivariate Taylor expansion recovers the joint probabilities that *j*_1_, …, *j*_*k*_ of the cohort reach stages 1, …, *k*.

### 2.3. Lifetime Reproduction PGF

For an individual born in stage *s*_0_, we define the *lifetime reproduction* PGF to encode the distribution for the numbers of offspring they produce from birth to age *y* + 1:

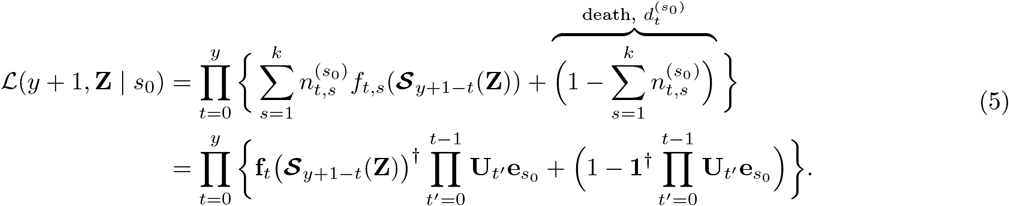

Above, the term **𝒮**_*y*+1−*t*_(**Z**) = (𝒮_*y*+1−*t*_(**Z** | 1), …, 𝒮_*y*+1−*t*_(**Z** | *k*)) in Eq (5) is a (column) vector of state distribution PGFs, with *s*-th entry the PGF for a single individual born to stage *s*. The (column) vector of stage-specific reproduction PGFs is denoted **f**_*t*_(***𝒮***_*y*+1−*t*_(**Z**)) = *f*_*t*,1_(***𝒮***_*y*+1−*t*_(**Z**)), …, *f*_*t,k*_(***𝒮***_*y*+1−*t*_(**Z**)). This vector has *s*-th entry the state distribution PGFs for a cohort of offspring born from a parent of stage *s*, denoted by *f*_*t,s*_ ***𝒮***_*y*+1−*t*_(**Z**)) = *f*_*t,s*_(𝒮_*y*+1−*t*_(**Z** | 1), …, 𝒮_*y*+1−*t*_(**Z** | *k*). The *s*′-th argument of *f*_*t,s*_(·) is *f*_*t,s*_(𝒮_*y*+1−*t*_(**Z** | *s*′), which describes the future state distributions for the numbers of the cohort born to stage *s*′ (from a parent of stage *s*).

Eq (5) tracks the future state of the sum of offspring produced by an individual over ages *t*, and up to that individual being aged *y* + 1 (and offspring up to age *y* + 1 − *t*). Uncertainty in the individual’s stage trajectory is found by probabilistically summing over stages the individual can be in at each age *t*, i.e.,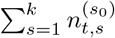. Given each stage *s* at age *t*, the PGF *f*_*t,s*_(·) yields the number of offspring the individual produces, born into different stages 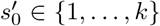 (matrix form: 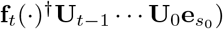. Offspring born when the individual was age *t* must live for *y* + 1 − *t* by the time the individual is aged *y* + 1. If offspring were born into stage 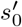, their current state distribution (at age *y* +1−*t*) is 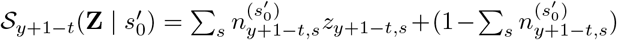. Given *f*_*t,s*_(*z*_1_, …, *z*_*k*_), a stage *s* individual’s reproduction PGF of offspring to stages 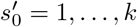 for each offspring stage 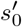 we replace the birth dummy variable 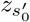 with the offspring’s current state, 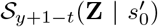. Substituting 𝒮_*y*+1−*t*_(**Z**) (the vector of survival PGFs) into *f*_*t,s*_ accordingly replaces newborns with their *current* (possibly dead) states. Lastly, offspring numbers produced at different ages *t* = 0, 1 …, *y* are independent random variables. The sum of independent random variables is given by the product of their PGFs.

To see that Eq (5) is a well-defined PGF, observe that for each age *t* and stage *s*, non-zero mortality means that 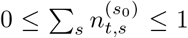. This means that the weighted sum 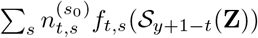 can result in an improper PGF. We correct this through the normalisation term, 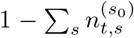 (the event that the individual has died). As such, setting all dummy variables to 1 results in 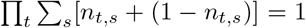. If the individual dies at age *t*′, then their contribution to their lifetime reproduction for all ages *t > t*′ is simply that they produce zero offspring with probability one.

To track the state of descendants across arbitrary generations *i* ∈ {1, 2, …,} within some fixed time *y*, we recursively compose the PGFs for lifetime reproduction. Let *t*_*j*_ represent ages at which the *j*-th generation descendants (*j < i*) produce the (*j* + 1)-th generation descendants. For the *i*-th generation descendants of an individual to be extant at the end of some time, *h*, we require that 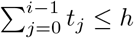. Then we recursively define

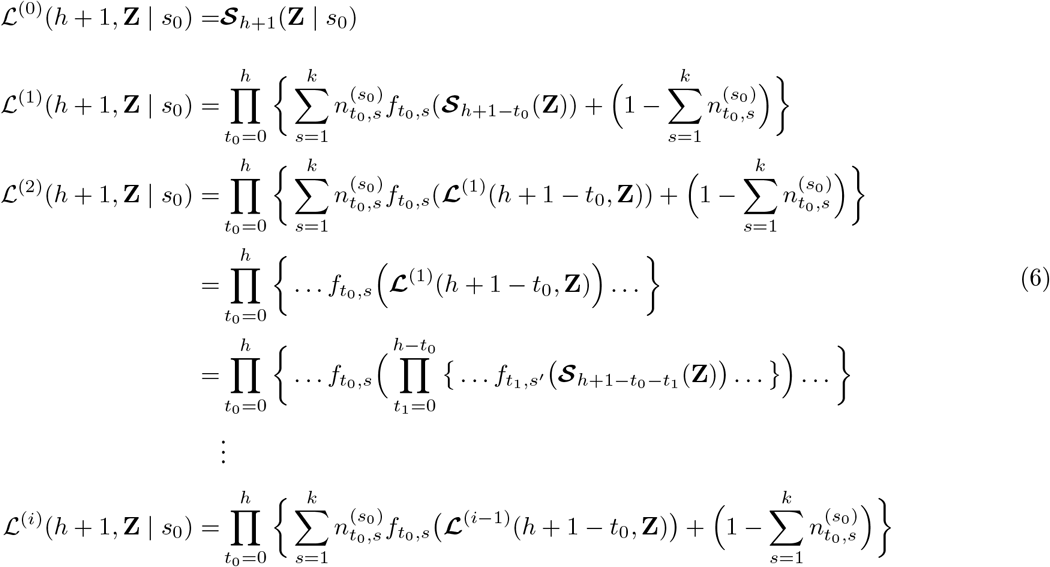

In the same way outlined after Eq (5), the matrix of dummy variables **Z** in Eq (6) allow the nested PGFs to track any generation of descendants who are at age *t* and stage *s*, while the individual which started the chain of descent is age *h* + 1. That is ℒ^(*i*)^ encodes the joint state distribution of the *i*-th generation descendants of an individual, accounting for all intermediate reproduction events which occur over (*i* − 1) generations, and within the time interval [0, *h* + 1]. In Box 1 we provide a graphical illustration of the lifetime reproduction PGF for a simple model with *t* = 0, …, 5 age-classes and *k* = 2 stages.

#### Box 1

**Example of the lifetime reproduction PGF**

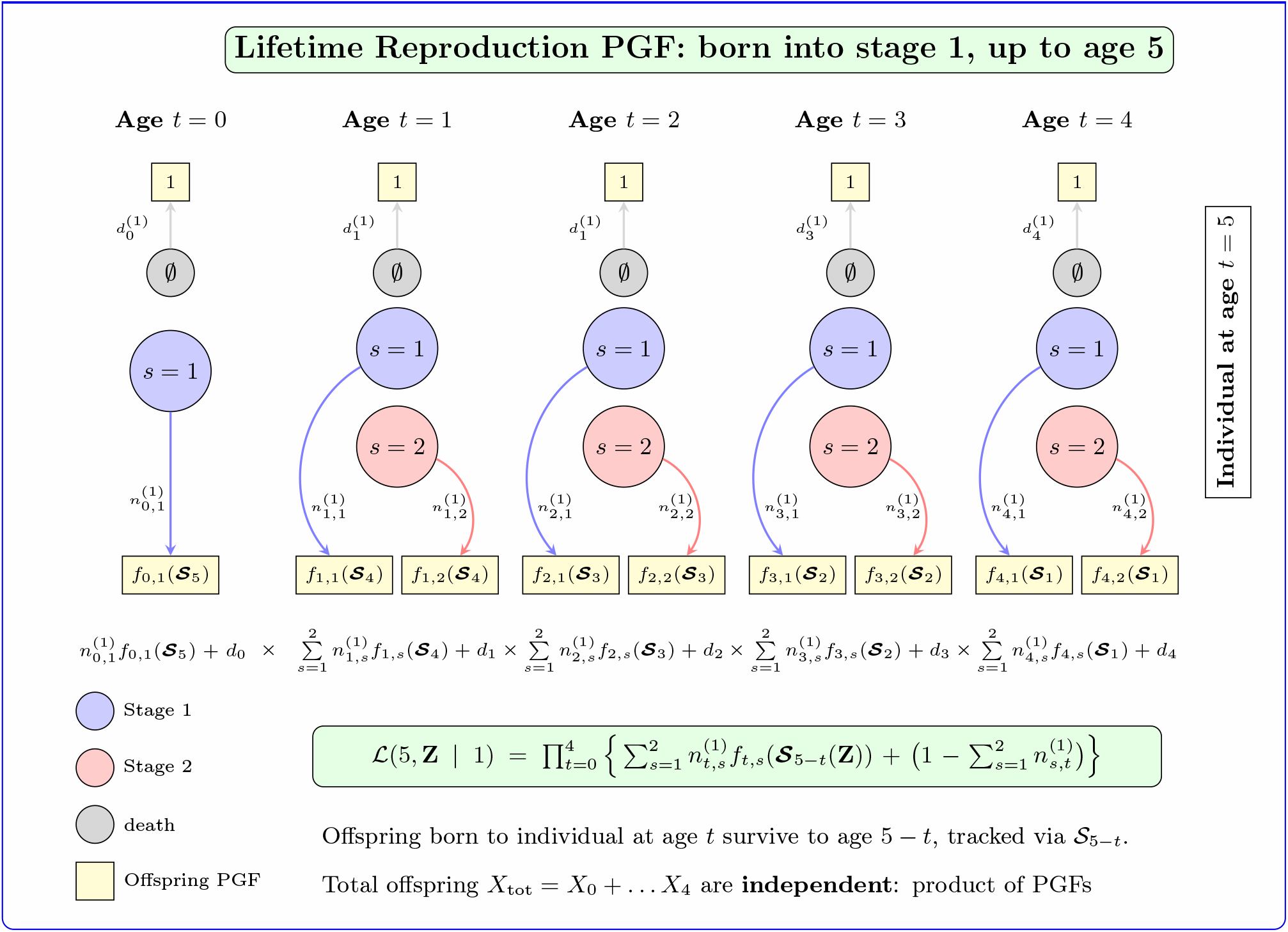

## 3. Kin formula

Suppose Focal is age *x* + 1 and born into stage *s*_*F*_ (we can relax this latter assumption). We define Focal’s kin through the [*g, q*] convention, introduced by Pullum [36], established by Coste *et al*. [1], Coste [21], and adopted by Butterick *et al*. [23] and Butterick [26]: kin related to Focal as the *g*-th generation descendant of Focal’s *q*-th ancestor. For the purposes of this model, we let *b*_*i*_ be the age of Focal’s *i*-th ancestor when producing Focal’s (*i* − 1)-th with *β*_*i*_ = *x* + 1 + ∑_*i*_ *b*_*i*_ defining the age of Focal’s *i*-th ancestor at present. We let the fertile ages be bound by 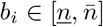.

In the next sections, we provide formulae for the joint PGFs for kin of Focal in three different regimes: descendants of Focal, collateral kin of Focal, and ancestors of Focal. We respectively denote these PGFs by Ψ^*g*^(**Z** | *x* + 1, *s*_*F*_), Ψ^*g,q*^(**Z** | *x* + 1, *s*_*F*_), Ψ^*q*^(**Z** | *x* + 1, *s*_*F*_). The PGFs are defined with arguments **Z** which is a matrix of dummy variables with entries *z*_*t,s*_ the numbers of kin of age *t* and stage *s*. Using the properties of generating functions we can then extract the joint probability of having 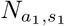 kin of age *a*_1_ and stage *s*_1_, and 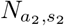 kin of age *a*_2_ and stage *s*_2_, and …, 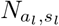 kin of age *a*_*l*_ and stage *s*_*l*_. We can equally extract the marginal probabilities of having 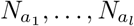 (or 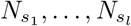) numbers of kin in ages *a*_1_, …, *a*_*l*_ (or stages *s*_1_, …, *s*_*l*_). In Appendix A we provide a brief recap in the algebraic extraction of such probabilities, and in Appendix F we show in practise how this is done numerically.

### 3.1. Descendants of Focal

Denote by *y*_*g*_ the age of Focal’s *g*-th generation descendant. Let 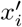 be the age at which the *i*-th generation descendant produced the (*i* + 1)-th generation descendant, with 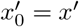 (the age of Focal’s reproduction). Then 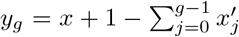. For Focal’s (*g* − 1)-th generation descendant to exist and produce Focal’s *g*-th, clearly we require 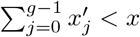. If this is satisfied, then the PGF for Focal’s *g*-th generation descendants is given by:

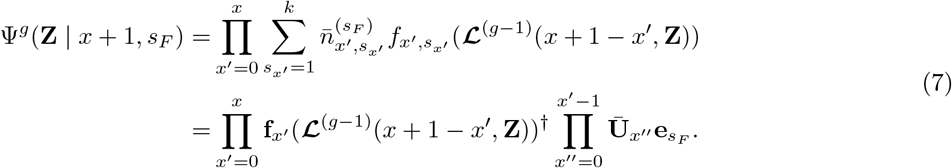

In Eq (7), 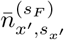 gives the state distribution of Focal at age *x*′ with 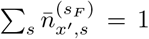 since Focal’s survival is guaranteed. Hence, there is no need for normalisation of the PGF for Focal’s lifetime reproduction. All of Focal’s descendants, however, are subject to mortality and have survival PGFs and lifetime reproductive output PGFs given through Eq (6). The term **f**_*x*_*′* ℒ^(*g*−1)^(*x* + 1 − *x*′, **Z**) = *f*_*x′*,1_ (ℒ^(*g*−1)^(*x* + 1 − *x*′, **Z**)), …, *f*_*x′,k*_ (ℒ^(*g*−1)^(*x* + 1 − *x*^′^, **Z**)) is a (column) vector of nested stage-specific reproduction PGFs, the *s* entry of which calculates reproduction over (*g* − 1) generations of descent starting from Focal’s offspring born into stage *s*. As per Eq (6), the above, through use of the matrix of dummy variables **Z**, keeps track of *g*-th generation descendants of Focal, over time horizon *x* + 1 − *x*^′^, and yields their probable age, stage and number. For each generation of descent *i < g*, we substitute the vectors of PGFs of lifetime reproduction for newborns into the newborn dummy matrix. Composing the PGFs for *i* = 0, 1, …, *g* − 1 procures the PGF for the children born to the (*g* − 1)-th generation: the *g*-th generation. For instance, regarding Focal’s grand-daughters (*g* = 2), we substitute ℒ^(1)^(*x*+1 − *x*^′^, **Z** | *s*) for the *offspring* dummy variables in the reproduction PGF for Focal: for every child of Focal born into stage *s*, we readily obtain the PGF for that child’s own descendants. A worked example is shown in Appendix C; Box 2.

### 3.2. Collateral kin

To obtain the PGFs for Focal’s collaterals, we need to know when Focal’s ancestor began the chain of events which results in the collateral’s current state distribution. For instance, Focal’s cousins descend through her aunts, who in turn descend from her grandmother, while Focal’s nieces descend through her sisters, who descend from her mother.

#### 3.2.1. Genealogical Markov Chains

To obtain the probable ages and stages of ancestral reproduction, We turn to genealogical Markov theory [37, 38]. So long as the multi-state projection matrix 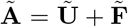 is irreducible^1^ there is a Markov chain with transition matrix **𝔓** = **𝔘** + **𝔉**. The entries of the transition matrix are defined through

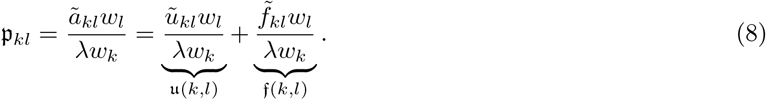

where **w** = [*w*_*i*_] is the stable population structure (the right Perron vector of **Ã w** = *λ***w**) and *λ* is the population growth rate (the dominant eigenvalue of **Ã**). We interpret *u*(*k, l*) as the probability that an individual currently in state *l* one-time step previously was extant and in state *k*. We interpret 𝔣 (*k, l*) as the probability that an individual was born from an ancestor of state *k* one-time step previously. Recall that state encompasses both age *t* and stage *s* through *kt* + *s*. That is, for instance, the proportion of individuals of age *t*^∗^ and stage *s*^∗^ is simply the (*kt*^∗^ + *s*^∗^)-th entry of **w**.

#### 3.2.2. Size-biasing

Recall that, given our definitions, 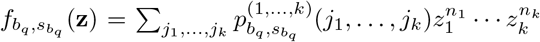 is the PGF for births by Focal’s direct *q*-th generation ancestor of state 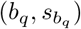. Suppose we want to find the PGF for sisters of Focal’s (*q* − 1)-th direct ancestor born in the same time-interval. That is, the other offspring of Focal’s *q*-th generation ancestor, excluding Focal’s (*q* − 1)-th ancestor. The probability that Focal’s (*q* − 1)-th ancestor is born into state 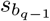 can be written

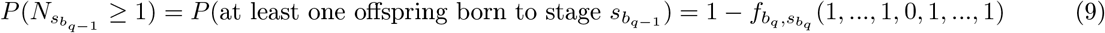

where the zero is in position 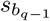. Condition on Focal’s (*q* − 1)-th ancestor being one of the 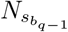 offspring born into stage 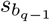. Let the numbers of other offspring born be random variables 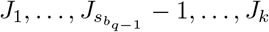, with probability:

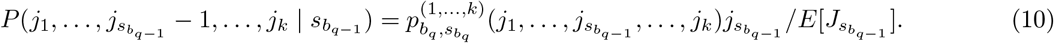

For notational ease, set 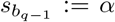. The PGF for the offspring excluding Focal’s (*q* − 1)-generation ancestor (who was born into stage 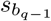), is found through the power expansion:

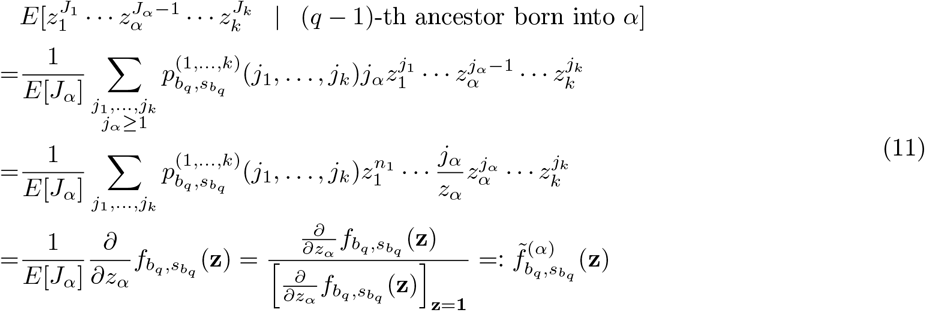

The size-biasing through 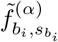, common in branching process models [39], removes one of ancestor’s offspring (Focal’s direct line) and provides the conditional distribution for the sisters of Focal’s direct line (additional of same age).

#### 3.2.3. The conditional PGF

If we condition on the event that Focal’s *q*-th ancestor produced Focal’s (*q* − 1)-th ancestor when aged *b*_*q*_ and of stage 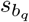, i.e., of state 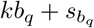, the (conditional) PGF for the state number distribution of Focal’s [*g, q*]-kin is:

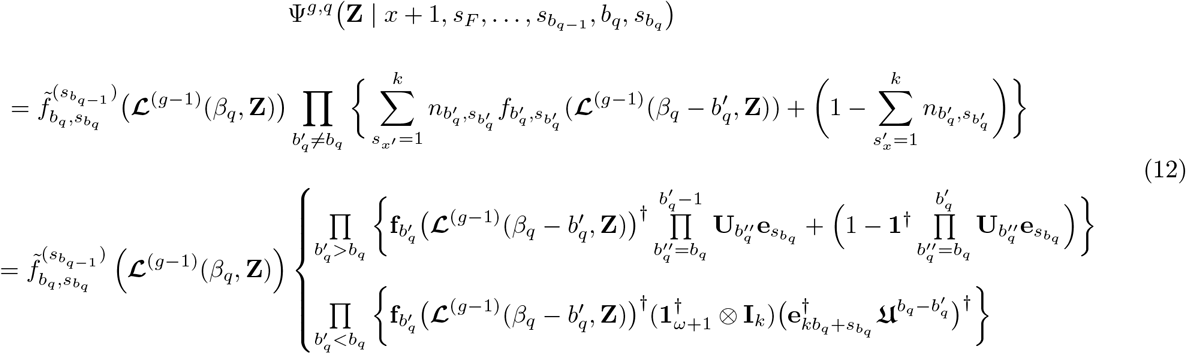

The bottom line of Eq (12) captures three mutually exclusive cases regarding the ages of reproduction of Focal’s *q*-th ancestor. The term to the left of the brackets covers reproduction of Focal’s *q*-th ancestor at age *b*_*q*_, when she produces Focal’s (*q* − 1)-th ancestor. The top line of the cases bracket accounts for reproduction of Focal’s *q*-th ancestor at ages 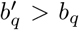, *after* Focal’s (*q* − 1)-th ancestor is born. The bottom line of the cases bracket accounts for reproduction of Focal’s *q*-th ancestor at ages 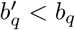, *before* Focal’s (*q* − 1)-th ancestor is born.

In more detail, consider the term to the LHS of the cases brackets. The composition yields the PGF for the lifetime reproductive output of the (*g* − 1)-th generation descendants of Focal’s *q*-th ancestor, whose line of descent is initiated by an offspring of Focal’s *q*-th ancestor born in the same time-interval as Focal’s (*q* − 1)-th ancestor. That is, kin who descend through one of Focal’s (*q* − 1)-th ancestor’s *same-age-sisters* [1, 23]. Recalling Eq (6), the term ℒ^(*g*−1)^(*β*_*q*_, **Z**) adds up the lifetime reproduction – depending on their survival – of (*g* − 1)-generations over the interval [0, *β*_*q*_], yielding the number age, and stage of the (*g* − 1)-th descendants. Inserting this PGF as argument of the size-biased PGF describing the reproduction of Focal’s *q*-th ancestor (in addition to Focal’s (*q* − 1)-th ancestor), 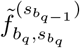, we obtain the PGF for the ancestor’s *g*-th generation descendants who descend through sisters of the child whose line leads to Focal.

Regarding the top line of the cases bracket, we know at age *b*_*q*_ Focal’s (*q* − 1)-th ancestor was born. For each age 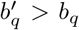 after this event, Focal’s *q*-th ancestor has transitioned stage and survived with probabilities in 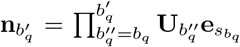 (i.e., the probability of being in stage *s* is the *s* entry). At each such age (if alive), and for each possible stage 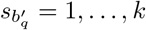 we calculate the PGF for her (*g* − 1)-th generation’s descendant’s life-time reproductive output. This is done by multiplying the (row) vector 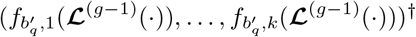 through the stage probability vector, 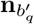, (conditional on survival). With probability 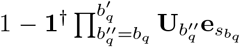 Focal’s *q*-th ancestor dies before age 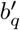 and has no offspring to form the chain of descent: this acts to normalise the sum of PGFs.

Regarding the bottom line of the cases bracket, we have conditioned on Focal’s *q*-th ancestor being age *b*_*q*_ and stage 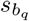 at birth of Focal’s (*q* − 1)-th ancestor. Therefore, when Focal’s (*q* − 1)-th ancestor is born we can write Focal’s *q*-th ancestor’s state distribution (in row vector form) by 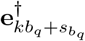. For each 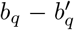 before the birth of Focal’s (*q* − 1)-th ancestor, the *q*-th ancestor had state distribution 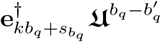. Her stage is found by applying 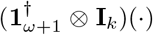. Pre-multiplying by 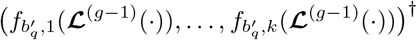 sums up the stage-specific PGFs for the (*g* − 1)-th generations lifetime reproductive output. No normalisation is required since we condition on *b*_*q*_ (Focal’s *q*-th ancestor must be extant for 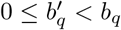).

#### 3.2.4. The unconditional PGF

Removing the conditioning, we face the problem that we don’t know the age of stage of Focal’s (*q* − 1)-th ancestor when she produced Focal’s (*q* − 1)-th ancestor. However we can probabilistically infer such quantities. From the genealogical Markov chain transition matrix, 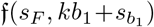 gives the probability that (newborn) Focal was born to mother of age *b*_1_ and stage 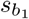 (where 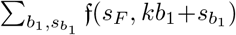 = 1). If mother was aged *b*_1_ and in stage 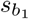 at Focal’s birth, the probability that at her own birth she was in state 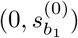 is given by 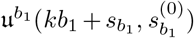 (where 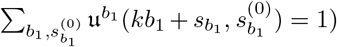. If mother was born to stage 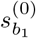, the probability that grandmother was in state (*b*_2_, 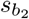) at this event is 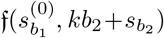 (where 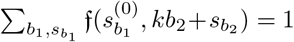). The conditional probability given *s*_*F*_, *b*_1_, 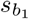, 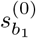, that gran had mother at age *b*_2_ and stage 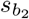 is 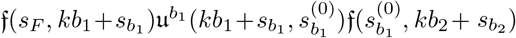 and the unconditional probability is 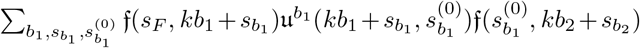 (extending the summation over 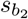 and *b*_2_ ensures the normalisation). As such, we can derive the conditional probability that Focal’s *q*-th ancestor procured her (*q* − 1)-th, given a specific chain of events defined by ancestral ages **b** = (*b*_1_, *b*_2_, …, *b*_*q*_), stages at reproduction 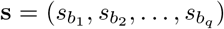, and stages at birth 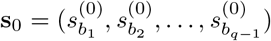. Let 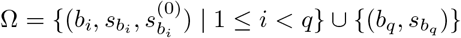. Let

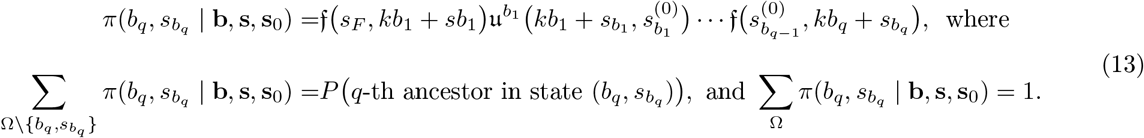

Then, the unconditional PGF becomes:

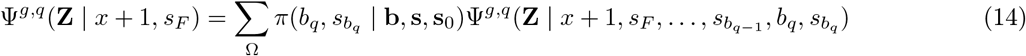

### 3.3. Ancestors

To construct PGFs for ancestors of Focal, we need to update Eq (2) to account for survival and stage transitions between two states in the life-course. Suppose that an individual is in stage *s*^′^ at age *t*_0_. We define the PGF describing their state at age *t > t*_0_, and accounting for survival, by:

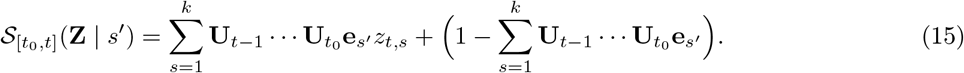

If we condition on Focal’s *q*-th ancestor being in state (*b*_*q*_, 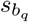) at reproduction of Focal’s (*q* − 1)-th ancestor, then the conditional PGF for the ancestor’s age and stage at Focal aged *x* + 1 is simply given by 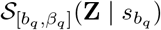. To obtain the unconditional PGF, similar to Section 3.2.4 we have to sum over all probable ancestral sequences which we might condition on:

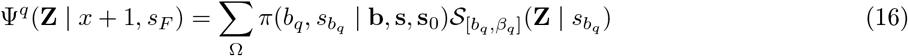

### 3.4. Including dependence on Focal’s stage distribution

Let **z** be a vector of dummy variable for Focal’s stage at age *x* + 1. If the kin-distribution for Focal’s kin is independent of the stage-transition path of Focal, as would be the case for collateral kin and ancestors, we see that the joint PGF for Focal’s kin and Focal are simply weighted sums of the PGFs defined respectively in Eq (14) and Eq (16), with weights given by Focal’s probable stage. That is, regarding collateral kin, we have

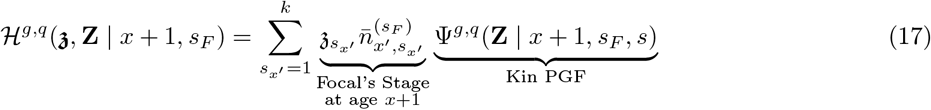

and regarding ancestor kin, we have

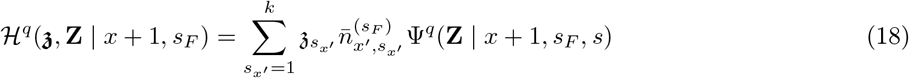

Alternatively, if the path of Focal explicitly defines the path of kin, as would be the case for descendants we use:

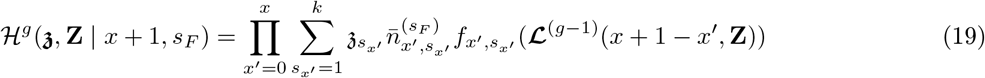

### 3.5. Counting dead kin

To count the number of kin which Focal loses, we update the kin state distribution in Eq (2) to include a dummy variable *d* which explicitly counts the death of an individual:

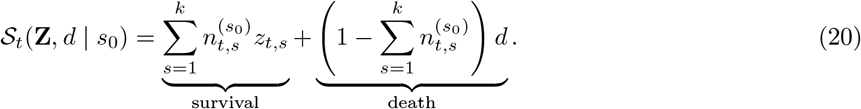

Recall the example following Eq (2), in which we consider a cohort of newborns 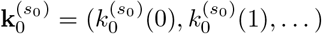 for *j* = 0, 1, …,, born to state *s*_0_. Suppose again we wish to know the probable numbers in stage *s* at time *t*. Substituting Eq (20) into the cohort newborn PGF 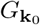, we now obtain the joint PGF for the number of survivors who end in stage *s*, and the number of deceased. For simplicity, again let 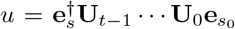 so that 𝒮_*t*_(*z*_*t,s*_, *d* | *s*_0_) = (*uz*_*t,s*_ + (1 − *u*)*d*). The we see

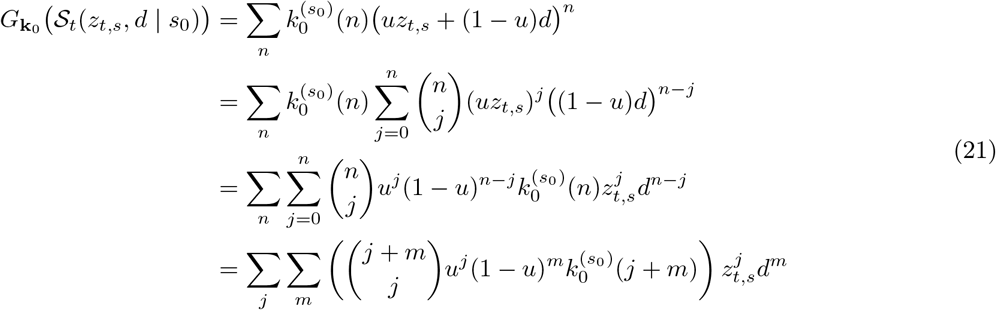

where in the last line we let the number of living of the cohort be represented by *j* and the number of dead by *m* = *n* − *j*. The joint probability that exactly *j* individuals are alive in stage *s* and exactly *m* individuals have died is then:

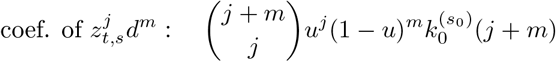

The updated state PGF in Eq (20) is particularly useful in analysing bereavement. For example, evaluating the coefficient at *j* = 0 for any *m >* 0 will yield the probability that none of the cohort are living at present, as a result of one or more dying. Furthermore, by substituting the full multivariate PGF 𝒮_*t*_(**Z**, *d*) into 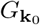 and taking the multi-variate expansion, one recovers the joint probability of the cohort which are living and in stages 1, …, *k* and the numbers of the cohort which have died.

#### 3.5.1. Lifetime reproduction PGF

In context of calculating the PGF for Focal’s kin, we need to update the lifetime reproduction PGF to now account for the reproduction of over generations, also counting the numbers who die in generation *i*. That is we define:

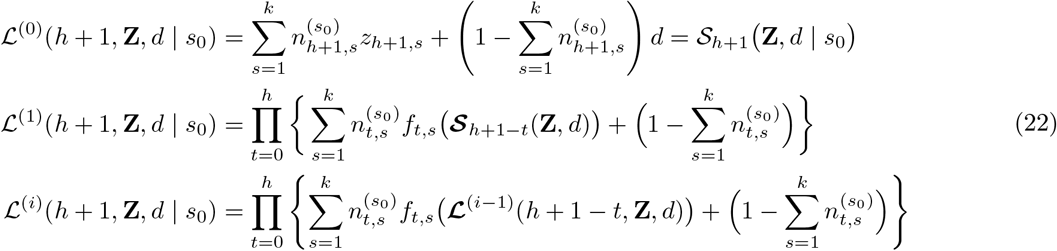

The above counts deaths in the *i*-th generation of descent, the recursion is structured such that the death dummy variable *d* is only active in the case ℒ^(0)^. This represents the descendants of interest. For all intermediate generations *j* ∈ {1, …, *i*}, the death of an ancestor results in the termination of that lineage, but does not add to the kin death count *d*

#### 3.5.2. PGF for descendants, counting kin loss

To update the PGF for Focal’s descendants Eq (7) and keep track of those of generation *g* which die, we simply substitute the updated kin state PGF, Eq (20), into the updated lifetime reproduction PGF, Eq (22). This approach ensures that if a descendant dies, their lineage stops (they reproduce no further), however, the kin are only tracked using the dummy variable *d* if of generation *g*:

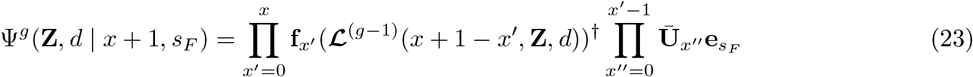

#### 3.5.3. PGF for Collateral kin, counting kin loss

To update the (conditional) PGF for Focal’s collateral kin, we must consider both deaths of the producer kin and all subsequent descendants which result in the lineages becoming extinct. We keep track of those kin which die only in generation *g* of descent from Focal’s *q*-th ancestor. The PGF for the *g*-th generation descendants of Focal’s *q*-th ancestor, counting the numbers which die, is:

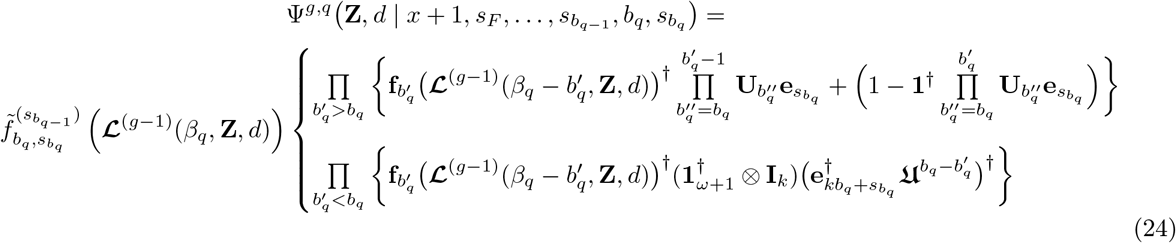

Consider the terms in the cases brackets. In the top line, the RHS term captures the case whereby Focal’s *q*-th ancestor dies before reaching age 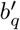. This event is not counted through the dummy variable *d* (we only count deaths of the *g*-th descendants of this ancestor). If the ancestor dies, this branch of the PGF stops (no further younger sisters of Focal’s (*q* − 1)-th ancestor are born). In the bottom line, we have conditioned on Focal’s *q*-th ancestor producing Focal’s (*q* − 1)-th at age *b*_*q*_. We know with certainty that she must be alive from birth to this age (we do not need to account for the survivorship of the ancestor; just their stage). In both cases 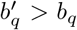 and 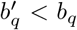, deaths of the subsequent *i* = 1, …, *g* − 1 generation descendants of Focal’s *q*-th ancestor are accounted for through the recursion **ℒ**^(*g*−1)^(·, **Z**, *d*). Only *after* generation (*g* − 1) has produced, however, do we count the number of deaths through the dummy variable *d* in the PGF **ℒ**^(0)^(·, **Z**, *d*) = *𝒮*_·_(·, **Z**, *d*). This ensures that we count the numbers of the newborns of the *g*-th generation descendants of Focal’s *q*-th ancestor which die between birth and Focal being age *x* + 1.

In the same way as in Eq (14), the unconditional PGF is found as a weighted sum of *π*(· | ·) over Ω of proper PGFs defined in Eq (24).

### 3.6. PGF for ancestors, counting kin loss

Here we consider the probabilities of an ancestor being alive in a specific stage versus being deceased. This depends on survival and stage transitions, e.g., an ancestor may start from stage *s*^′^ at age *t*_0_ and end at age *t > t*_0_ when Focal is *x* + 1. We also track death of the ancestor. As such, using the dummy variable *d* to count death, we define:

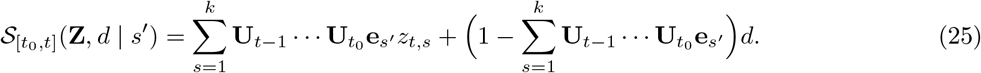

Similar to Eq (16) of Section 3.3, the (conditional) PGF for Focal’s *q*-th ancestor, counting whether the ancestor is dead, is 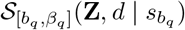. The unconditional PGF is found as a weighted sum of the conditional PGFs, over the ancestor probability space Ω:

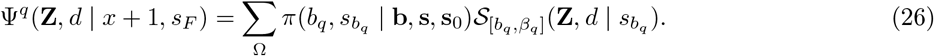

Some examples of how the above PGFs yield the probable numbers of kin-loss are given in Appendix C.2.

## 4. Application

Here, we use parity as a demographic stage which structures kin-networks. We use four stages *s* = 1, 2, 3, 4 respectively defined through parity classes 0,1,2, and 3+. We use UK parity specific fertility rates from 1964-2023, sourced from the ONS [40], and projected for the 2025 period using the model of Ellison *et al*. [41]. We use age-specific mortality rates from 1964-2023, sourced from the ONS [42], and projected using the model of Hilton *et al*. [43]. We don’t assume that parity level affects mortality (and thus the same age-specific mortality schedules applies to all parity levels). In the parity stage model, newborns start life in the first age-class and are of parity 0. Survivorship follows a Bernoulli distribution. We assume that parity-and-age-specific offspring number follow Poisson distributions. In Appendix G, as extra, we show how the model reproduces the expected numbers of kin found by the parity-structured model of Caswell [18].

### 4.1. Offspring

In Fig 1 we plot the marginal stage distributions for Focal’s offspring by age of Focal. As illustrated, the probability Focal has non-zero daughters (i.e., at least one daughter of any parity level) is zero up until she is age 15, at which point she enters her reproductive life. After age 15, the probability that Focal has one or more daughters increases with the offspring strictly being of parity 0 up to Focal being around age 40 (when we see the possibility of her daughters having their own offspring and moving to parity 1).

**Figure 1.**
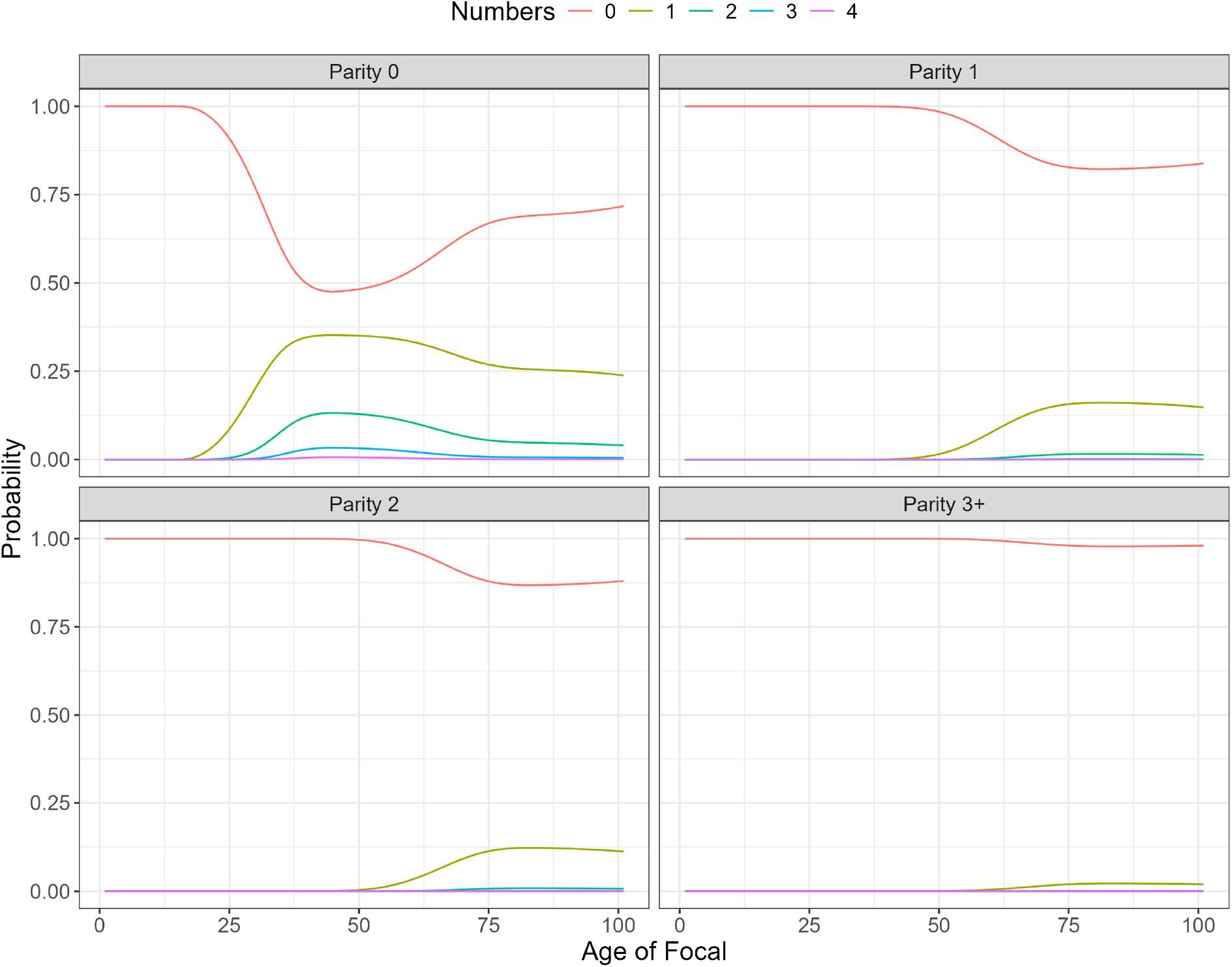
The marginal parity number-distributions of Focal’s daughters, by age of Focal.

In addition to the marginal stage distributions, the model can extract the joint parity distributions. The probability that Focal has *N*_1_ daughters in stage 1 (parity 0), *N*_2_ in stage 2 (parity 1), *N*_3_ in stage 3 (parity 2), and *N*_4_ in stage 4 (parity 3+). In Fig 2 we plot the ten most probable occurrences of such joint number-stage distributions, for a range of ages over Focal’s life-course.

**Figure 2.**
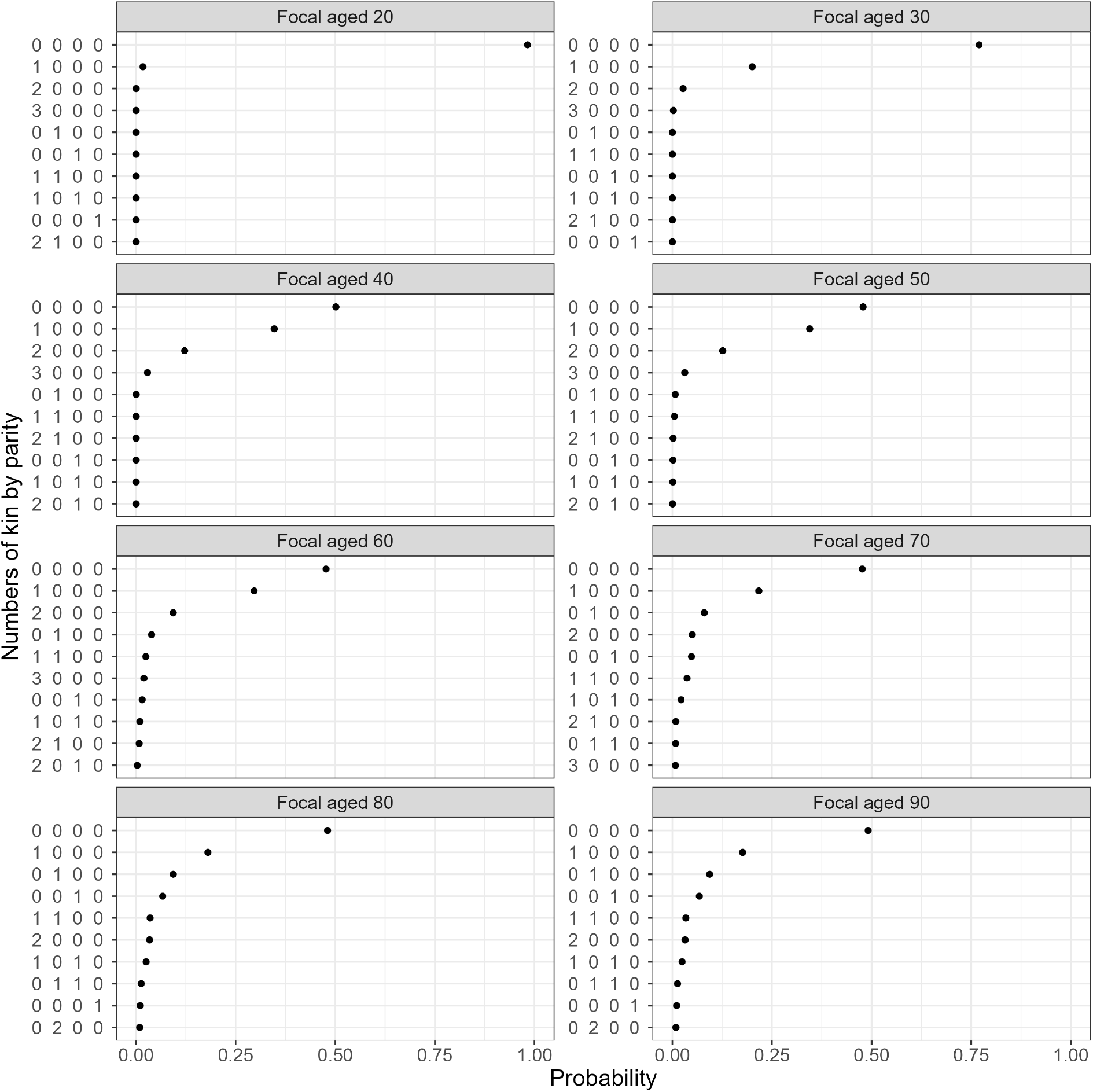
Joint distributions of the number of daughters of Focal, by age of Focal and parity of daughter. The strings *N*_1_, …, *N*_4_ along the *y*-axis represent (joint) events of Focal having: *N*_1_ daughters of parity 0, *N*_2_ daughters of parity 1, *N*_3_ daughters of parity 2, and *N*_4_ daughters of parity 3+. For each age of Focal, we plot the 10 most probable.

### 4.2. Sisters

Using the same methods as the case for daughters, the marginal parity number-distributions for sisters. As shown in Fig 3, we see that when Focal is born there is just over a 0.25 probability that she has one or more sisters (upper-left panel). The probability that she has one or more sisters of parity 1,2, or 3+, however is negligible (upper-right, lower-left and lower-right panels). By the time Focal is 15, there is about a 0.48 probability that she has one or more sisters in parity 0 (upper-left panel). The probability that Focal has exactly one sister of parity 1 is maximum when Focal is around age 50.

**Figure 3.**
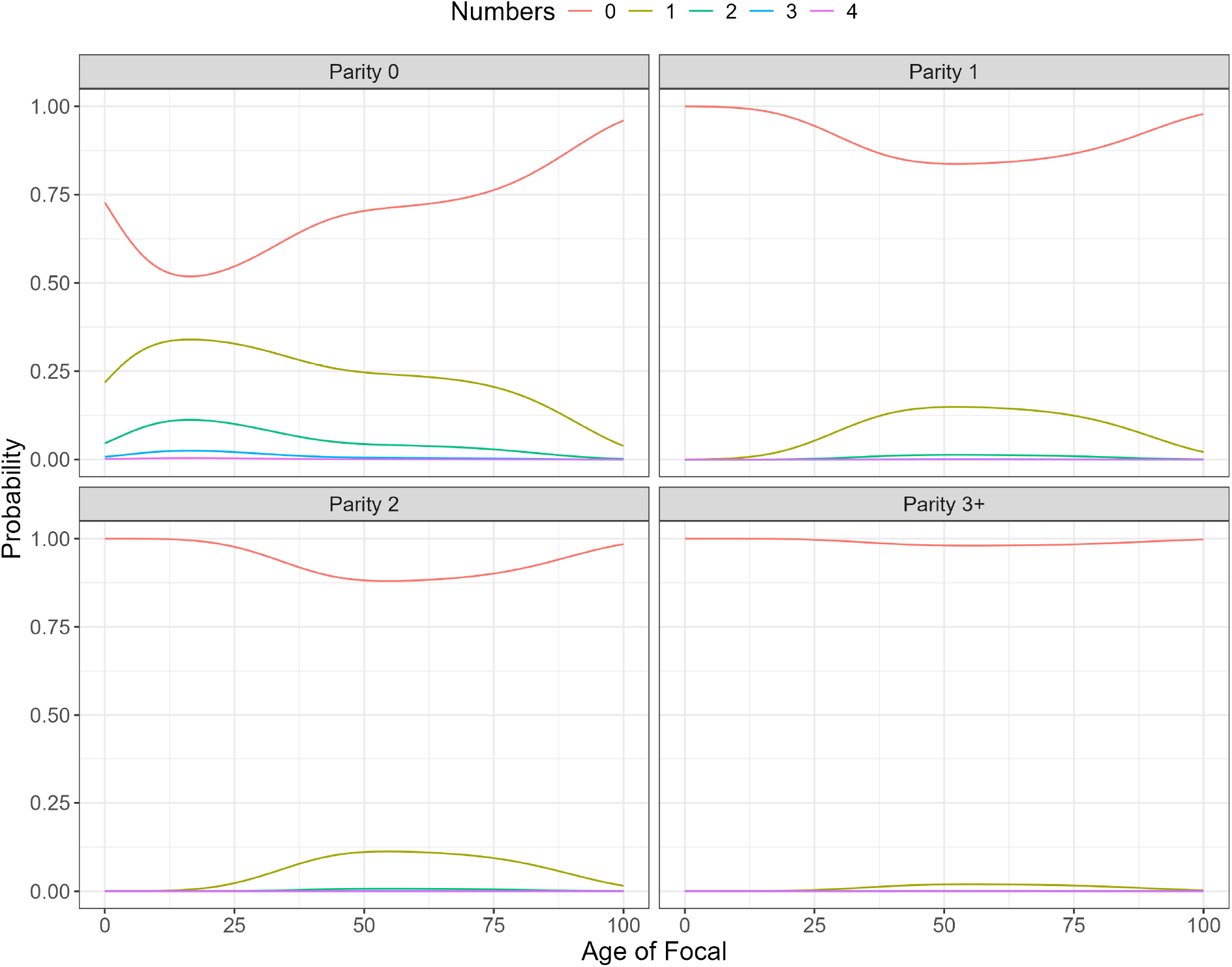
The marginal parity number-distributions of Focal’s sisters, by age of Focal.

In Fig 4 we plot the probabilities that Focal has *N*_1_ sisters in stage 1 (parity 0), *N*_2_ in stage 2 (parity 1), *N*_3_ in stage 3 (parity 2), and *N*_4_ in stage 4 (parity 3+). Due to the one-sex population modelling, for all ages of Focal, the most probable number-parity combination is that Focal has no sisters (zero of parities 0,1,2, or 3+), and the second most likely that she has one sister in parity 0 and no other. The third most probable outcome changes however. As Focal ages from 30 to 40, we see that it becomes more likely that Focal has one sister in parity 2 (and no other sisters) compared to her having 2 sisters in parity 0 (and no other).

**Figure 4.**
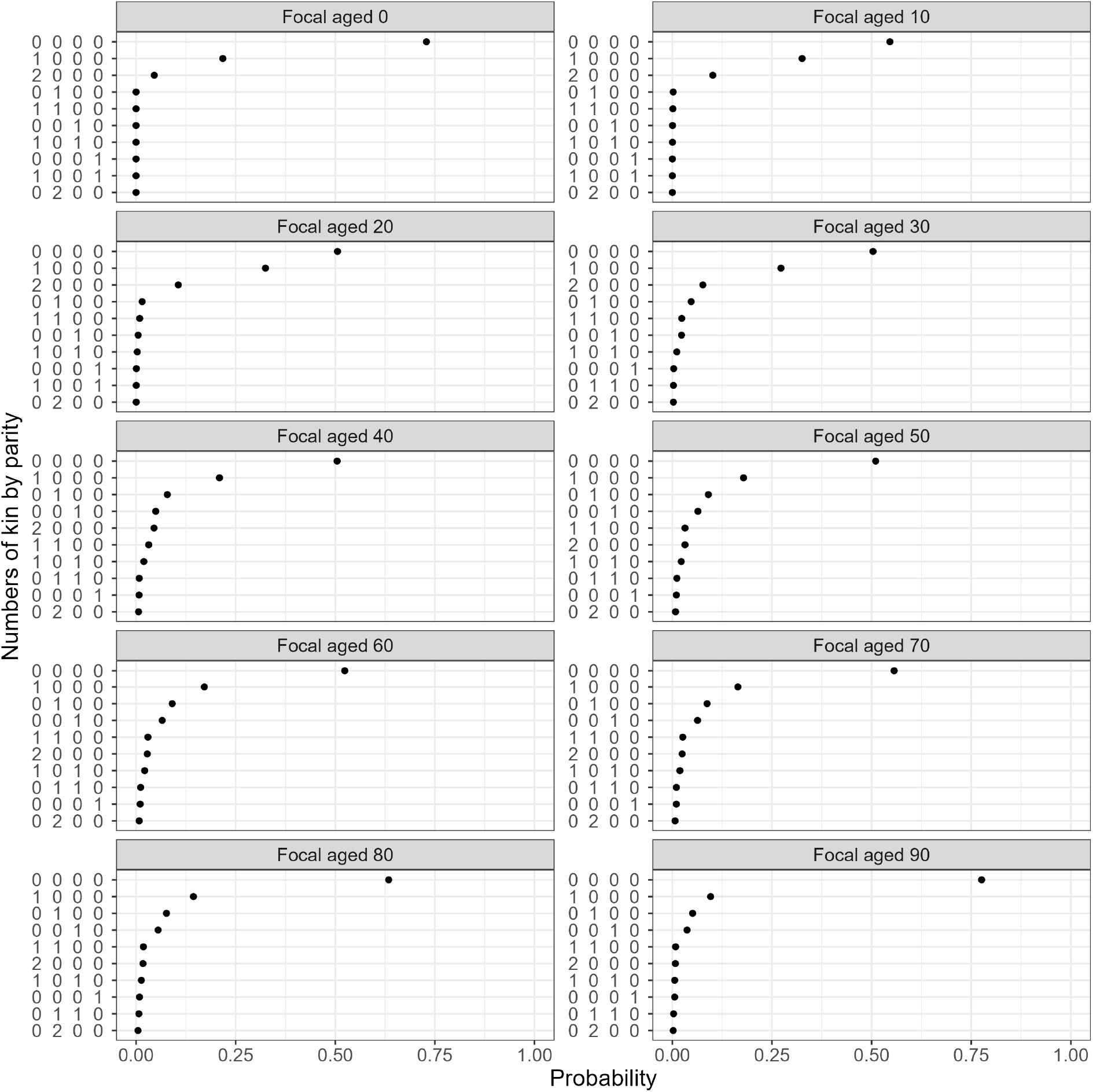
Joint distributions of the number of sisters of Focal, by age of Focal. The strings *N*_1_, …, *N*_4_ along the *y*-axis represent the (joint) events of Focal having: *N*_1_ sisters of parity 0, *N*_1_ sisters of parity 1, *N*_3_ sisters of parity 2, and *N*_4_ sisters of parity 3+. For each age of Focal we plot the 10 most probable.

### 4.3. Conditioning on Focal’s parity: the childless cohorts

Using Section 3.4 we calculate the joint probabilities that Focal and kin are in given states. For instance, consider the case in which Focal is childless but has at least one sister – a demographic phenomenon characteristic of the 1960s cohort in the UK [44]. The probability that Focal is in parity 0 at age *x*+1 is given by 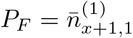. The joint probability that Focal is in parity 0 at age *x* + 1 and has *N*_1_ = 0, …, *N*_4_ = 0 sisters in parity-levels 0, 1, 2, and 3+ (i.e., zero sisters) is given by 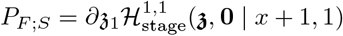. As such, the probability that Focal is childless but has one or more sisters is given by 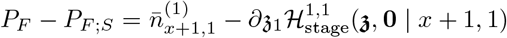. In Fig 5 we compare, for a typical Focal individuals of different ages, sampled from the population at different periods in time, the probabilities that they are childless but have one or more sisters.

**Figure 5.**
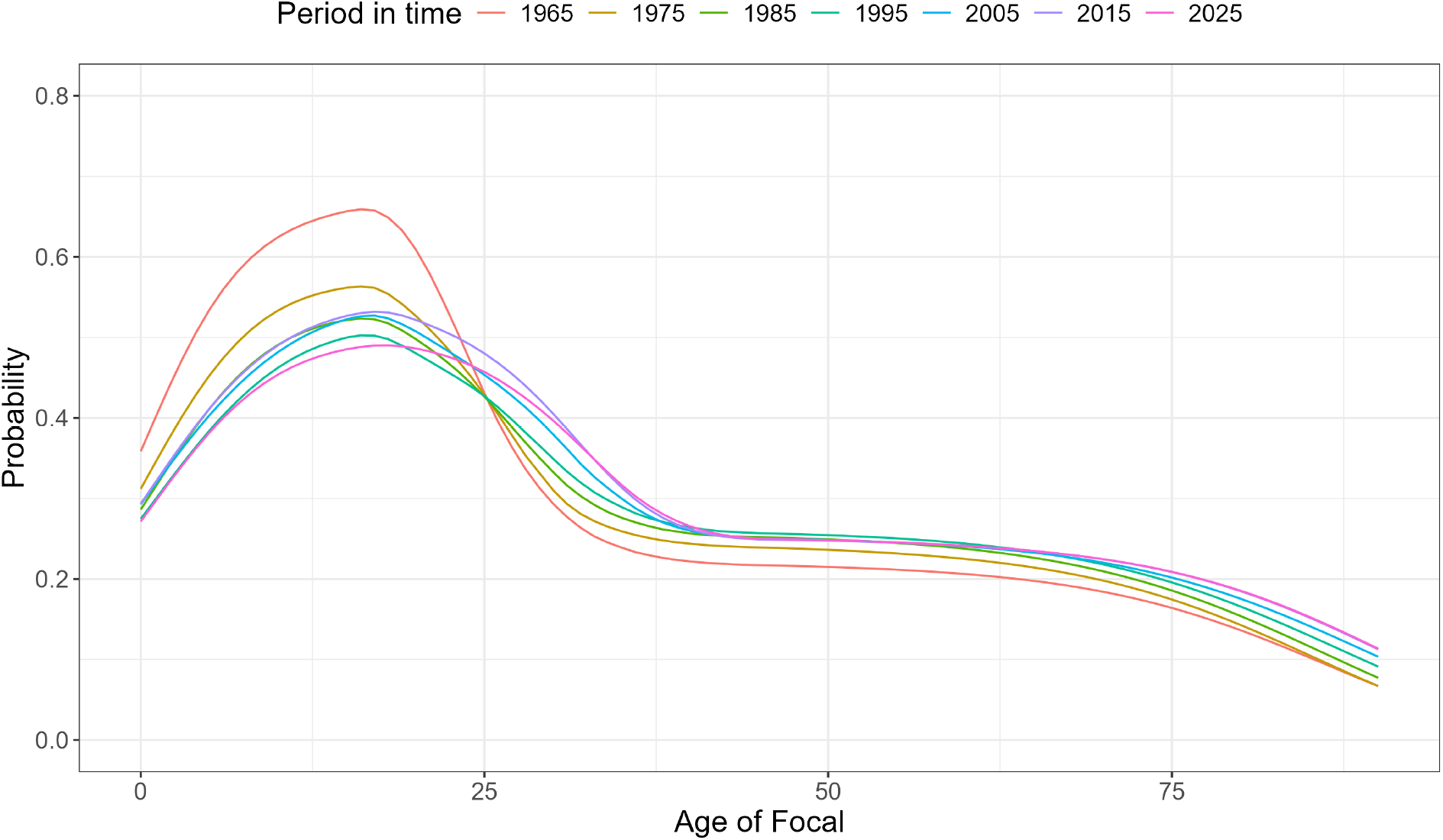
The probabilities that Focal has no children but at least one sister, by age of Focal. We compare period based predictions.

### 4.4. Bereavement of kin

Using Section 6, here we consider the PGF for the number of Focal’s kin including those which have died, Ψ(**Z**, *d* | *x* + 1, *s*_*F*_). By setting **Z** = **1** we obtain the marginal PGF for kin which Focal has lost between her being born and age *x* + 1. In Fig 6 we respectively plot the probabilities that Focal loses *j* = 0, 1, …, daughters or sisters, by age of Focal, and as a function of the period in which we sample Focal.

**Figure 6.**
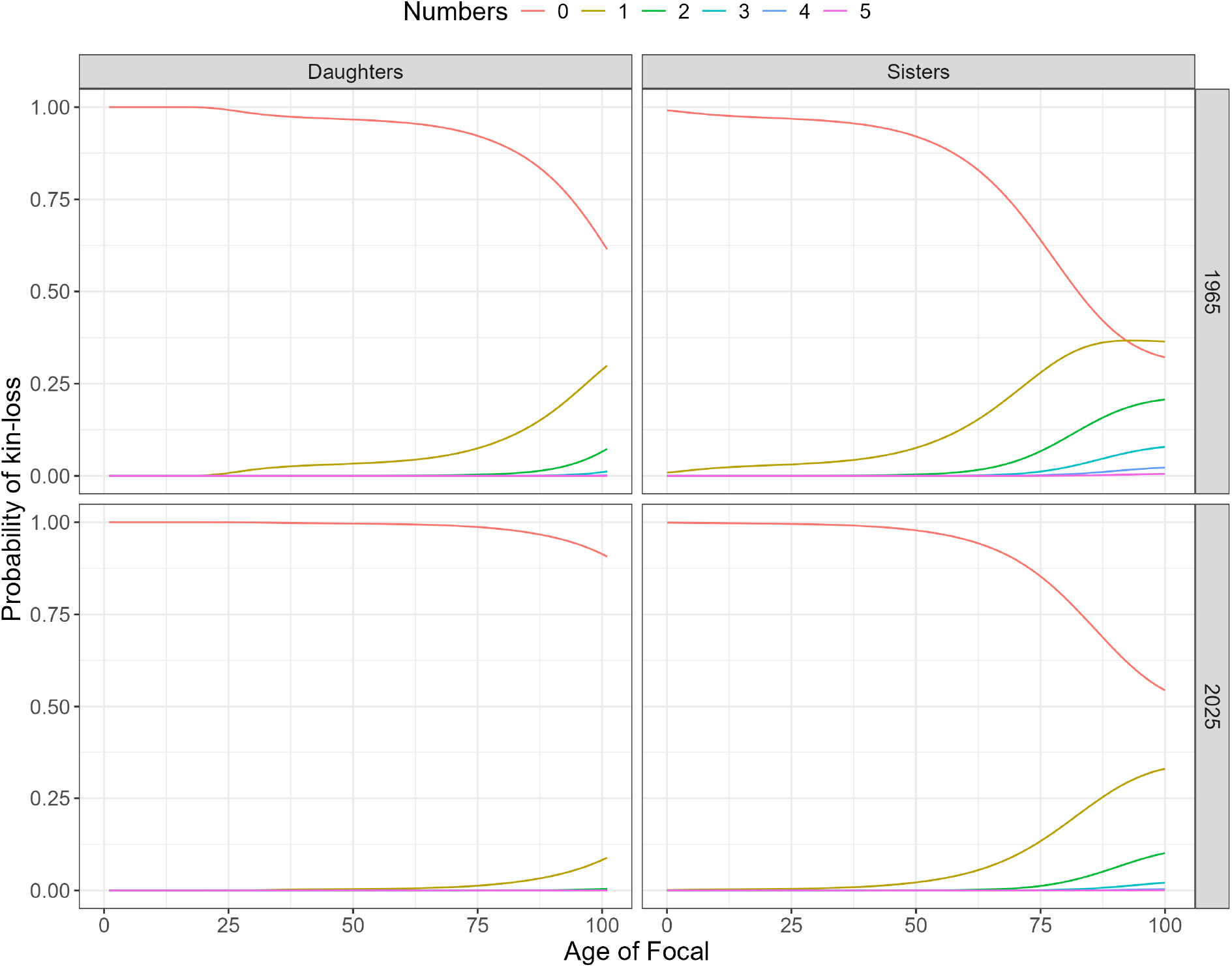
The probabilities that Focal loses so many children and sisters, by age of Focal. Note that these probabilities include the probability that the kin were never born.

The state of Focal being kinless – an important topic in human demography which has attracted considerable empirical attention [45, 46] – is also possible. In our framework, being kinless corresponds to the extinction of a specific lineage. Suppose for instance, that Focal at age *x* + 1 has no kin as a result of *one or more of the kin dying* between Focal’s birth and her present age. The probabilities that *m* = 1, 2, … kin deaths result in Focal having no living kin are found through 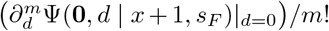. In Fig 7 we illustrate these probabilities (where *j >* 4 kin death probabilities are omitted for visual ease) regarding the cases of Focal’s daughters and sisters. Note the different scales on the *y*-axis.

**Figure 7.**
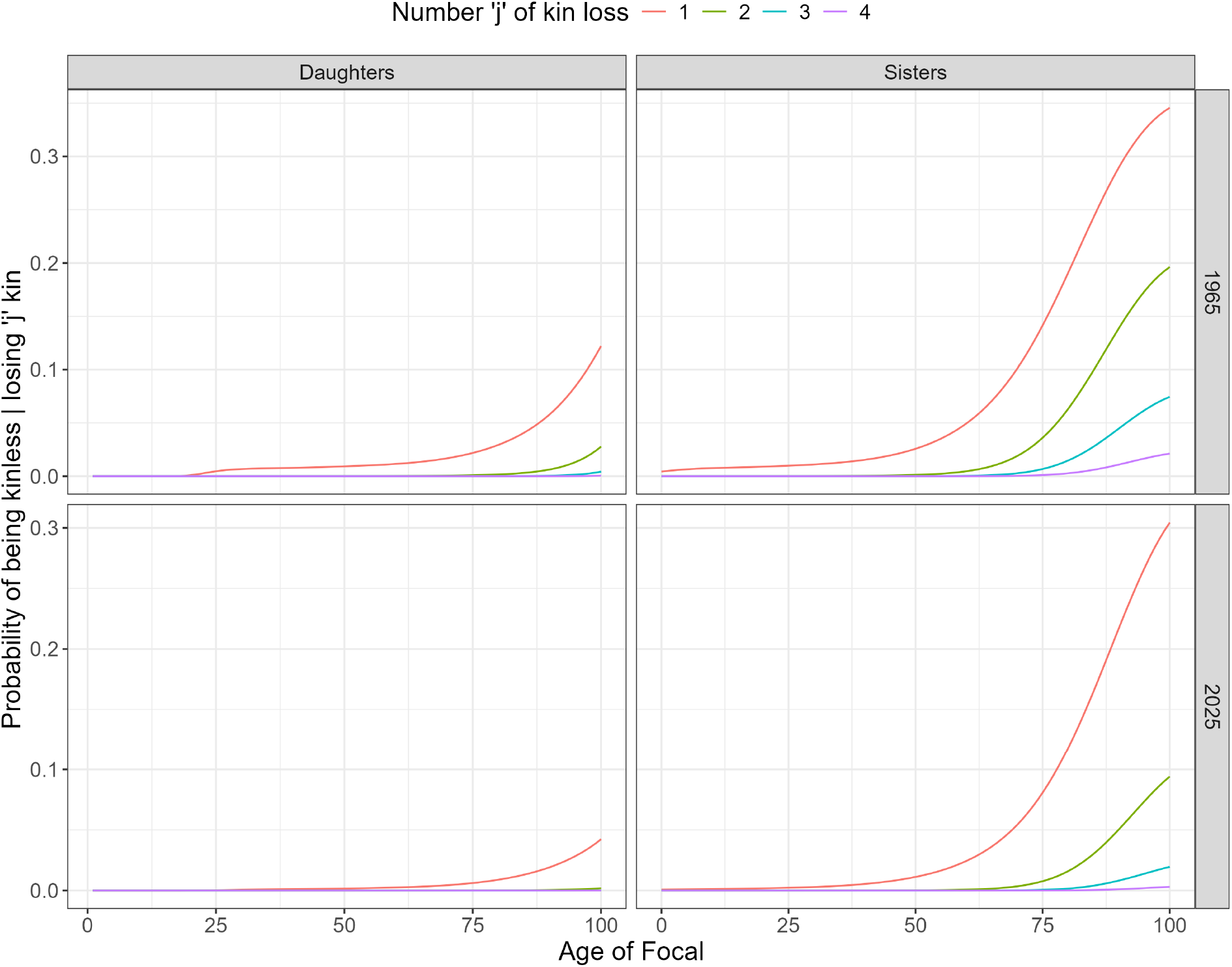
The probability that Focal has no kin-type due to *j* = 1, 2, 3, 4 of that kin-type dying. Left; daughters, right; sisters. We compare the 1965 period based prediction to the 2025 period.

As a final demonstration of the model output, we consider the probable numbers of Focal’s granddaughters which are orphaned. A derivation of how to apply the PGF modelling approach to extract such events is given in Appendix E. Here, in Fig 8 we plot the results for the likeliness that one or two (higher numbers have negligible probability) of Focal’s granddaughters are orphaned, by age of Focal. Comparing a Focal who lives her life under the 1965 period rates and one who experiences the 2025 rates, we see a considerable difference in the probabilities. Under the 1965 rates, by age 95 we find a 13% chance Focal has an orphaned granddaughter. Under the 2025 rates we find a 4% chance.

**Figure 8.**
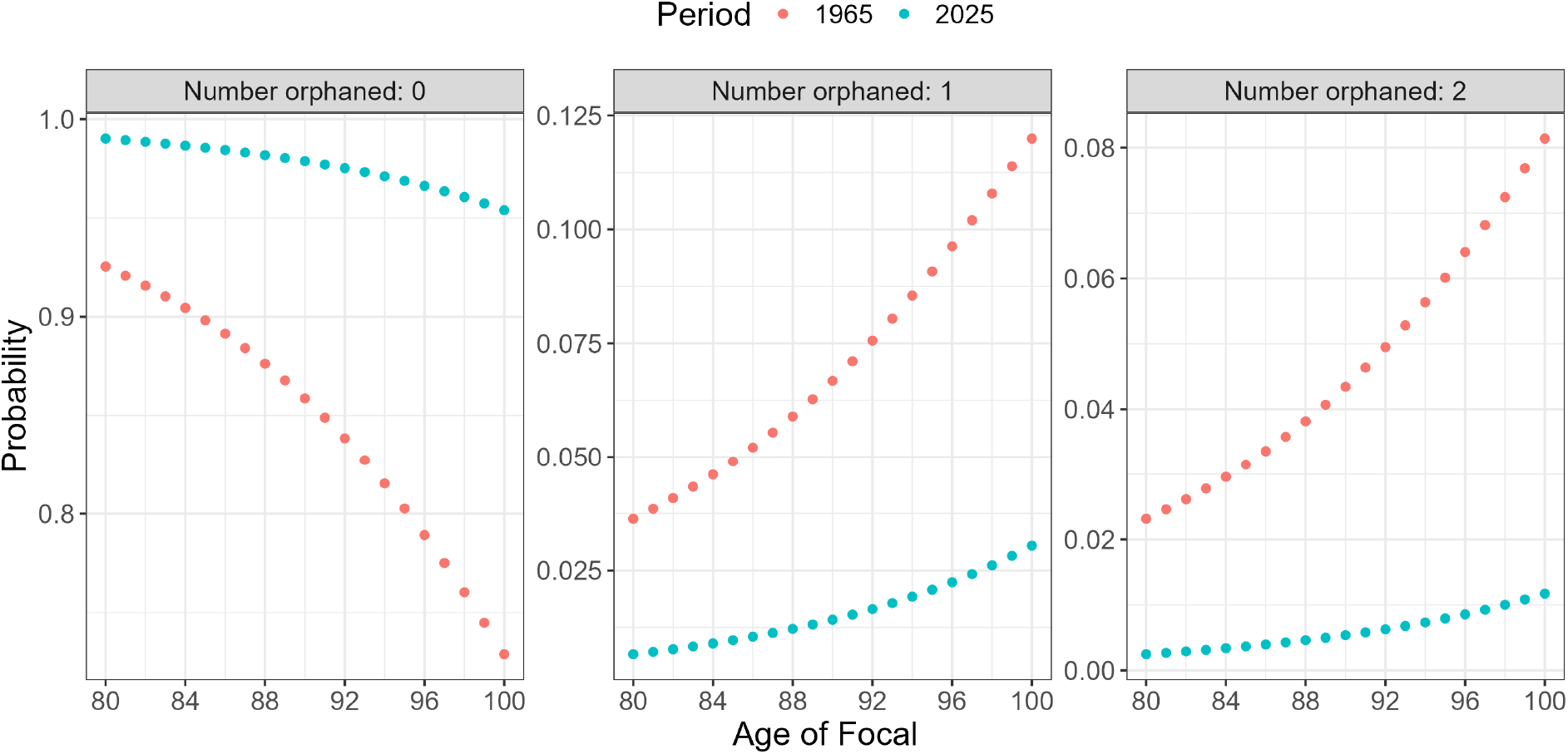
The probable number of Focal’s granddaughters who have lost their mother. We compare the 1965 period based probabilities to the 2025 period.

## 5. Discussion

This research presents the first analytical model to predict the probable numbers of kin in an age×stage structured population. In principal, our method allows one to calculate (i): the probabilities that an individual has so many kin of different ages *and* in different stages. (ii): the probabilities that an individual has so many kin in different stages (or ages) when marginalising over age (or stage). (iii): the marginal number distributions (PMFs) for kin structured only by stage or age. Extensions of the model show (iv): how to calculate the joint probability that an individual is in a given state (age×stage) *and* the joint probabilities of having so many kin of different ages and or stages (see Section 3.4), and (v): the probabilities that an individual loses so many kin-types, where kin are structured by generation number (see Section 6). In addition to generation number, extensions could include the age or stage of death of kin. Knowing the age of kin-loss, for instance, allows for appropriate support to bereaved parents [47].

The framework herein provides a comprehensive and novel probabilistic analysis of the kin-network. In humans, as well as the parity example, stages cover a range of demographic characteristics which people transition between. The joint distribution for Focal’s and her kin’s stage can define health needs [48], educational attainment [49], financial support [50], and emotional care [51], but to name some. We use parity in the application as a simple illustration of the models use. Nevertheless, motivated by the knowledge of the 1960s UK cohort on average having fewer children but more siblings, we show how likely it is that Focal will be childless but have more than one sister. Ongoing research suggests that 20% of the 1960s cohort are likely to be childless, with implications for kin available to care [9]. Our model provides a fast way to calculate the exposure (population size) for such a cohort of childless Focals who have sisters, circumventing the need to run complex and time-demanding micro-simulations such as Soc-Sim [52]. In animal populations, the numbers of an individual’s kin which have dispersed [53, 54] or which have died [55] are well-known to affect reproductive success. For instance offspring which delay leaving a group might benefit from group protection and at the same time help raise others’ offspring [56]. Additionally, dependencies on maternal figures from early age can affect individual mortality schedules [57], and thereby, also affect a population’s growth rate [2].

We demonstrate population distributions for the numbers of kin by age and stage of both Focal and her kin, and for arbitrary kin types. This approach allows for analytical results regarding demographic uncertainty that approach the complexity of those obtained via microsimulation. Our model requires as inputs, probability distributions for age-and-stage-specific mortality and fertility and stage transition probabilities. Expanding the rate distributions through power series naturally produces the probability generating functions from which the kinship formulae are derived. One has the flexibility to choose from empirical or parametric distributions. Parametric have the advantage of potential analytic tractability (although at the expense of demographic realism), while empirical have the advantage of being more suitable for the population of interest (however will require numerical methods when deriving the PGFs – see Appendix F).

Mathematically, we apply theory of branching processes to the field of kinship [22, 23]. Our model is an alternative representation of a Crump-Mode-Jagers branching process (e.g., see [39]), similar to a Multi-type branching process with discrete types composed of states (*a, t*): the age *a* and stage *s* of Focal’s kin. Focal and Focal’s kin’s states change respectively according to the transition matrices **Ū** and **U**. These linear algebra operators emulate the stochastic movement between age, stage, and death in state space (just and and stage for Focal who is immortal). The recursive nature of the branching process PGFs come in the form of the *f*_*a,s*_(·) which describe the *offspring distribution* for any kin of state (*a, s*). The recursive relationship between PGFs of different generations ensure that if Φ^(*g*)^(*z*) is the PGF for generation *g*, then Φ^(*g*)^(*z*) = Φ^(*g*−1)^(**f** (*z*)). The recursive nature of ℒ^(*i*)^ (see Eq (6)) allows one to nest ℒ^(*i*−1)^ inside the fertility function *f*_*t,s*_ of ℒ^(*i*)^. In essence, this function composition defines a multi-type branching process.

There are some caveats regarding the assumptions we require for applying to branching processes. First, independence of reproduction *between individuals* – a defining characteristic of a branching process – and *at different ages* for one individual, is not always realistic in natural populations [58]. In humans for instance, elapsed time between births can and may even be correlated with inherited behaviours [59]. The model is also time-homogenous, restricting it’s ability to represent kin-distributions in real cohorts for which age and time increment in tandem. A careful indexing with time-variant rates and a time-inhomogeneous genealogical Markov chain (e.g., like the one proposed by Butterick *et al*. [60]) would be all required to extend this framework to time-variant rates. Lastly, at present our model lacks a male population. Including an additional dimension of sex would result in block-structured vectors of the kin-state PGFs: 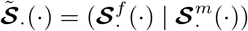 where 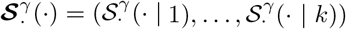 for individual of sex *γ* = *f, m*. State-specific reproduction PGFs would also be structured by sex, e.g., the vector of reproduction PGFs for an individual of sex *γ* would be 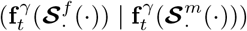. The *s*-th argument of this vector, 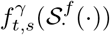, would yield the PGF describing the state of a cohort of female offspring born to the parent in stage *s* (with *s*^′^-th argument the number of this cohort born into stage *s*^′^). The (*k* + *s* +1)-th argument 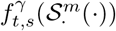, would yield the PGF describing the state of a cohort of male offspring born to the parent in stage *s*. Incorporating two-sexes should not be theoretically demanding, although would increase the computational complexity of the model.

In this research we propose a methodology that, for the first time, enables the derivation of complete joint probability distributions for the number of an individual’s kin, structured by age and stage. Such richness of output goes beyond existing approaches – mostly confined to expected values. Our model opens the door to ask questions of how probable it is that an individual has a given arrangement of living or dead relatives at any point in their life. For living kin, we show what demographic characteristics define them and how the kin’s state may be related to the individual. Our model jointly tracks how many kin within each generation die, and the possibility of orphanhood. The research here will prove important in understanding the availability of kin-based support networks and informal care in humans. The framework will have implications for research on population ecology, with in particular emphasis on changes in relatedness in state-structured animal populations.

## Appendix A

**Using the PGFs to extract probable kin numbers**

Recall that we denote age×stage state by (*t, s*). Given a matrix valued kin PGF Ψ(**Z** | ·), we can find the joint probabilities for the numbers 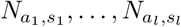 of kin in states (*a*_1_, *s*_1_), …, (*a*_*l*_, *s*_*l*_), by applying the multi-variate Taylor expansion:

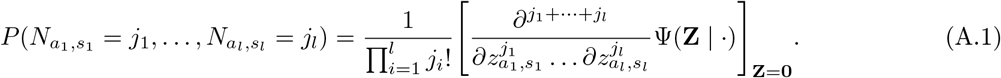

Respectively marginalising over stages or ages we recover vector valued PGFs of kin, structured respectively by age *a* (regardless of stage), or stage *s* (regardless of age). The vector valued PGF for kin-numbers structured by age is found by setting *z*_*a,s*_ = *z*_*a*_ for all *s*; the vector valued PGF for kin-numbers structured by stage is found by setting *z*_*a,s*_ = *z*_*s*_ for all *a*:

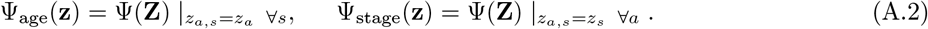

From the marginal vector-valued PGFs, respectively, we find the joint probabilities that Focal has 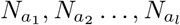 kin of ages *a*_1_, …, *a*_*l*_, and the joint probabilities that Focal has 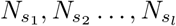 kin in stages *s*_1_, …, *s*_*l*_. These probabilities are found through Taylor expansions:

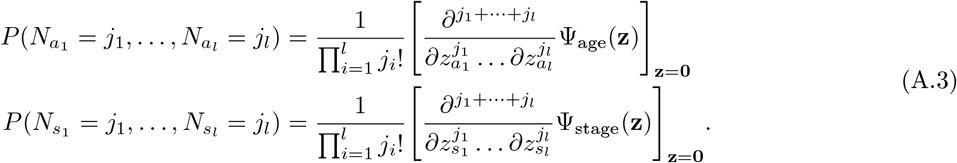

The marginal age PMF for the numbers of kin of age *a*_*i*_ (marginalising over all other ages and all stages) is found by setting 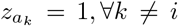 in Ψ_age_ (i.e., we recover the scalar PGF 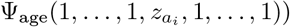. The marginal stage PMF for the numbers of kin in stage *s*_*i*_ (marginalising over all other stages and all ages) is found by setting 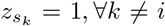. in Ψ_stage_. The respective probabilities, 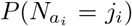: that that *j*_*i*_ kin are of age *a*_*i*_, and *P* (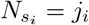: that *j*_*i*_ kin are in stage *s*_*i*_, are found using

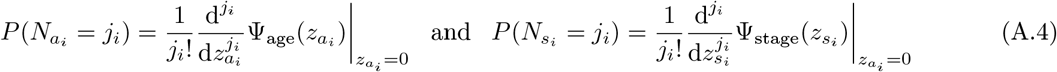

Lastly, marginalising over ages and stages, we recover the total numbers of kin (unstructured). The total numbers of kin are found by setting *z*_*a,s*_ = *z* and taking the univariate Taylor expansions.

## Appendix B.

**Relationship to Butterick *et al*. [23]**

In an age-structured demography, the matrices **U**_*x*_ become scalars *u*_*x*_ and the reproduction PGF becomes *f*_*x*_(*z*) = ∑_*j*_ *P* (*X* = *j*)*z*^*j*^. Consider older sisters which are of age *s*_1_. Using notation of Butterick *et al*. condition on the event that mother had Focal at age *b*_1_:

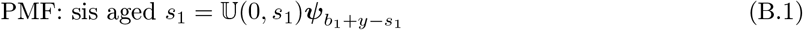

where the *i, j* entries of 𝕌 (0, *s*_1_) are 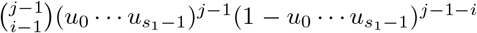, providing probabilities that *i* − 1 out of *j* − 1 individuals (offspring of mother) survive from age 0 to age *s*_1_, and entries 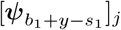 provide probabilities mother has *j* − 1 offspring for *j* ≤ *Q*. Letting 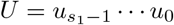, in the present framework we have:

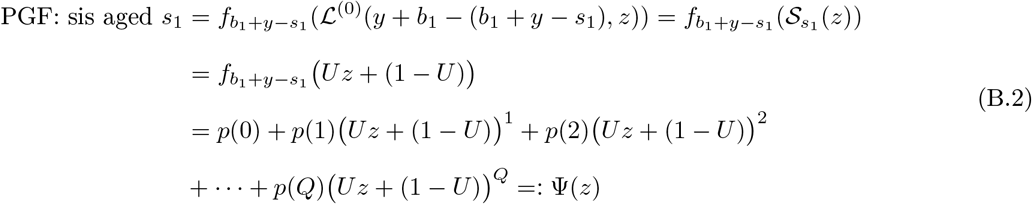

Consider first, the probability that Focal has one sister. The model of Butterick *et al*. calculates this probability as the second row of U multiplied through ***ψ***:

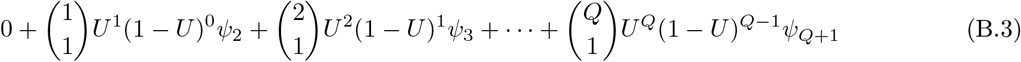

which becomes:

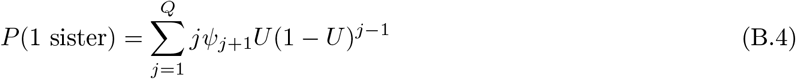

Regarding the PGF model, observing that *p*(*i*) = *ψ*_*i*+1_, we see Eq (B.2) reproduces the same result:

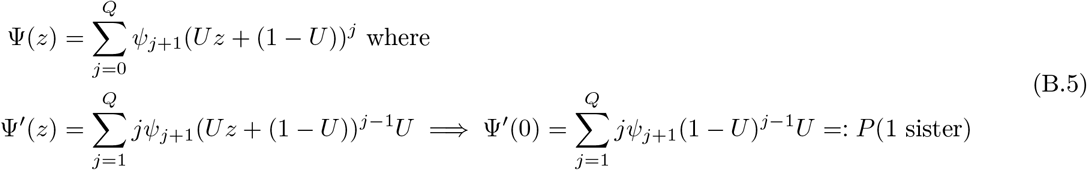

For general *k*, Butterick *et al*. [23] find probability that Focal has *k* sisters as the (*k* + 1)-th row of the matrix 𝕌 multiplied through ***ψ***:

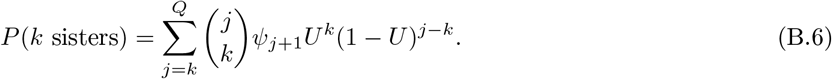

In the present framework, using the PGF Ψ(*z*) defined in B.2, we find the exact same probability of having exactly *k* sisters, albeit using the *k*-th coefficient in the Taylor expansion evaluated at *z* = 0:

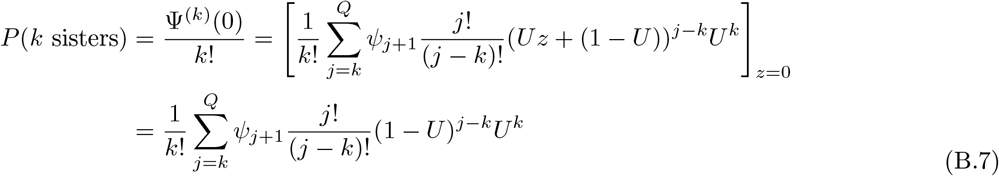

whence dividing through *k*!:

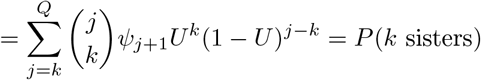

Over all ages the sister could be, because we know the sum of random variables 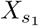 for *s*_1_ = 1, …, *n* with measures given by 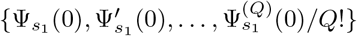 is the convolution of its measures, we recover Eq (34) in Butterick *et al*. [23].

In more detail it is possible to show formally, that the two frameworks are equivalent. We do so through proving that the survival and fertility components of the models are the same.

### Appendix B.1. Survival

Here we show that 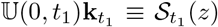. Suppose *Q* is chosen such that 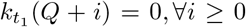 Let again 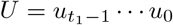. Regarding the PMF model, we know that the (*j* + 1)-th row of 𝕌 (0, *t*_1_), written [𝕌 (0, *t*_1_)]_*j*+1_ provides the probability that *j* kin survive:

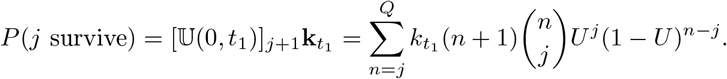

Let the PGF be

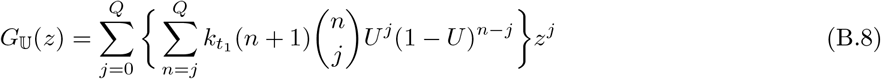

Regarding the PGF model, the probability that *j* kin survive is found by taking the *j*-th Taylor coefficient in the expansion of the PGF 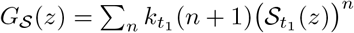 where 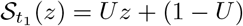.

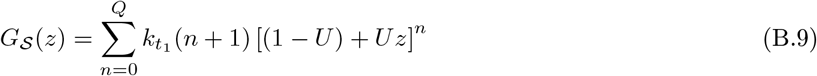

where

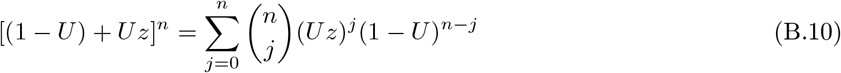

which substituted back results in:

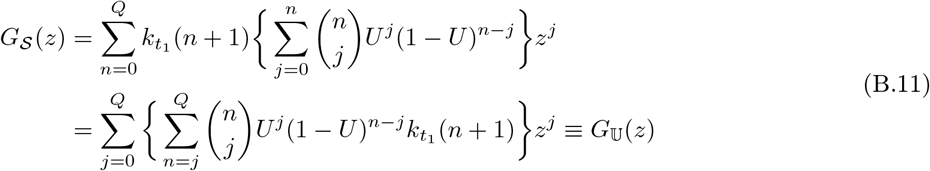

Comparing coefficients for each *j* we further see

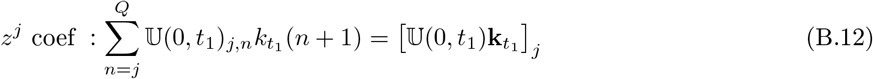

### Appendix B.2. Reproduction

In the framework of Butterick *et al*. [23], the number-distribution of offspring born to kin of age *s* (represented by a number PMF **k**_*s*_) is 𝔽 (*s*)**k**_*s*_, where the matrix 𝔽 (*s*) has *l*-th column entries given by elements of the vector of the *l*-th power convolution of ***ψ***_*s*_. That is,

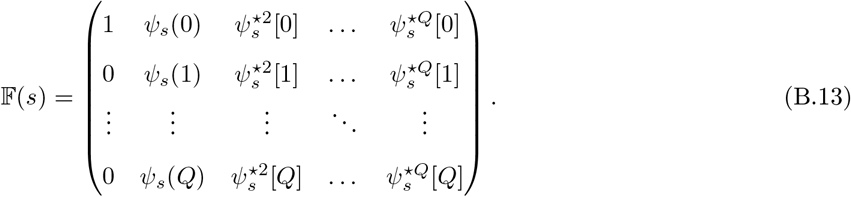

In the present framework, with no stages, the reproduction PGF is 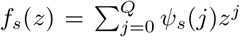 (i.e., the PGF for the number of offspring born to an individual at age *s*). The PGF for the number of offspring born from *l* independent individuals is the *l*-fold convolution:

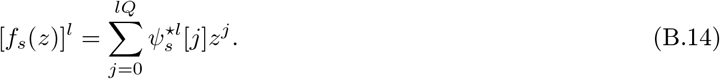

The methods are identical because the *l*-th power in the PGF [*f*_*s*_(*z*)]^*l*^ corresponds precisely to the (*l* − 1)-th column (the *l*-th convolution) of the 𝔽 (*s*) matrix: 𝔽 (*s*) is simply the linear algebra representation of the composition operator on the probability space. For instance, suppose that we have a distribution of “aunts” of age *s* represented by a vector **a**_*s*_, the product 𝔽 (*s*)**a**_*s*_ yields a vector whose entries give the probable numbers of newborn cousins to aunts of age *s*, and equally, are the coefficients of Ψ_*A*_(*f*_*s*_(*z*)) where Ψ_*A*_(*z*) is the PGF for number of aunts.

More generally, let **off** be the PMF for the number of offspring kin, and **pro**_*s*_ be the PMF for producer kin (at age *s*), so that **off** = 𝔽 (*s*)**pro**_*s*_. Let *G*_**off**_(*z*) be the PGF for offspring number. Since

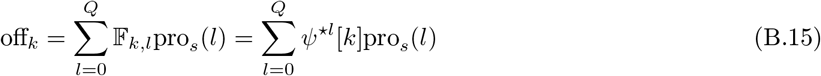

the PGF is

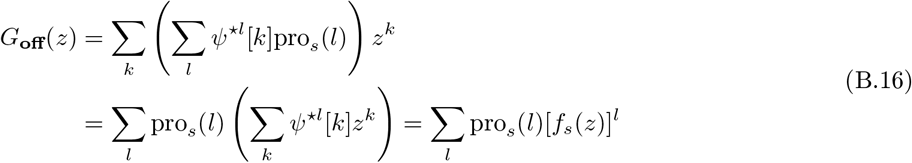

which is the composition of PGFs

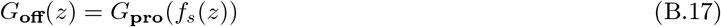

## Appendix C

**Examples**

### Appendix C.1. Examples of the PGF model

#### Box 2

PGF for grand-daughters

The PGF for grand-daughters is found using Eq (7). We substitute the lifetime reproduction PGFs for daughters into the offspring variables for Focal’s reproduction. For instance, if Focal reproduces over ages *x*′ = 0, 1, …, *x* then we have:

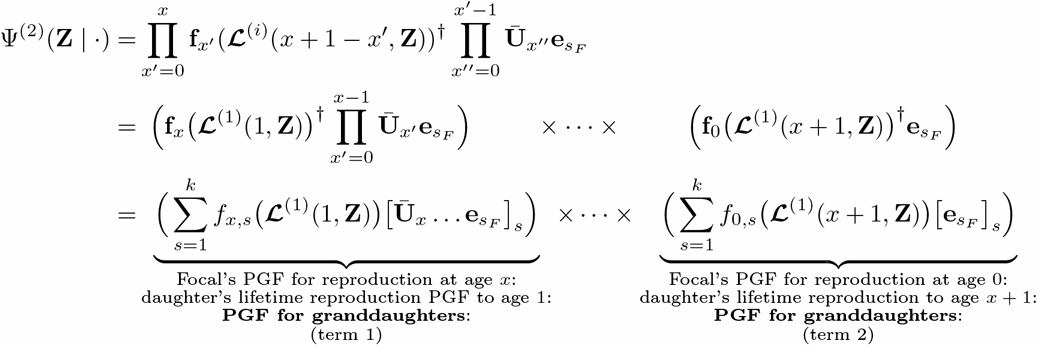

Expanding “term 1” and “term 2” we see:

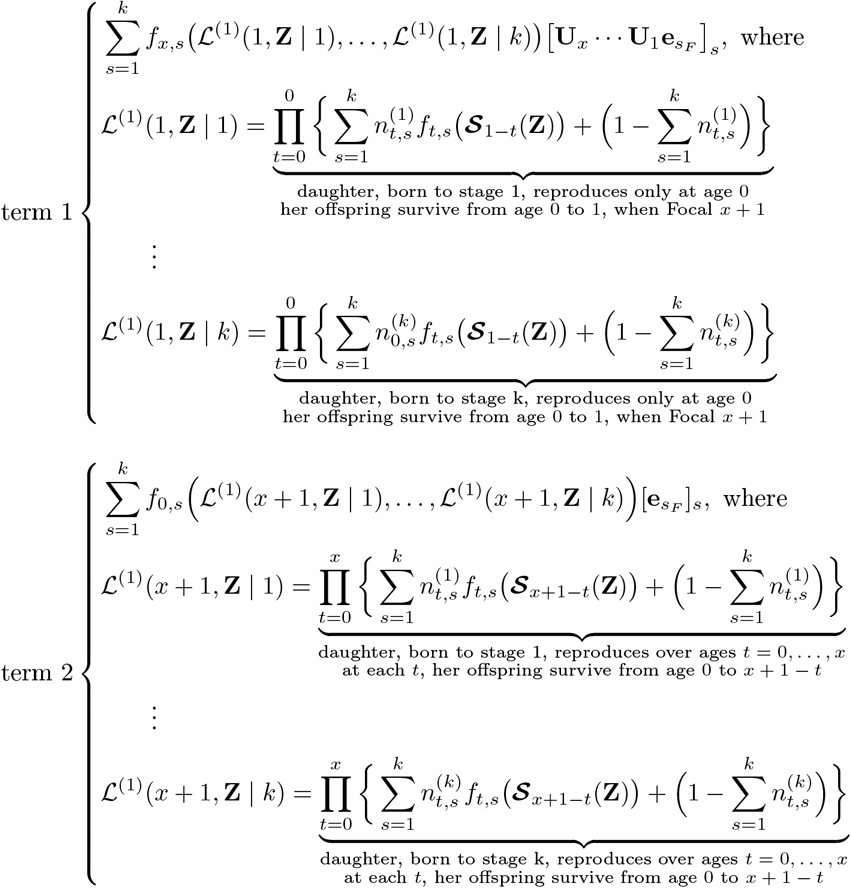

### Appendix C.2. Example of the PGF model with death

#### Box 3

PGF for daughters

In this simplified example, consider 2 stages *s* = 1, 2, two ages *t* = 0, 1, Focal enters life into stage 1 with numbers defined by the fertility PGF **f** (*z*). Suppose that Focal is now at age 1 and consider the joint probability that Focal has *j*_1_ daughter in stage 1, *j*_2_ daughter in stage 2, and *m* daughters who have died. The probability daughter survives and is in stage *i* = 1, 2 is *U*_*i*_ = [**U**_0_*e*_1_]_*i*_. The probability that daughter dies is 1 − *U*_1_ − *U*_2_. The kin state PGF accounting for death becomes:

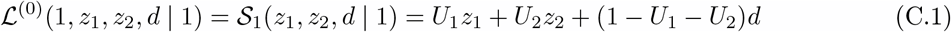

which we plug into the fertility PGF and multiply through ages of Focal’s reproduction (here only at age 

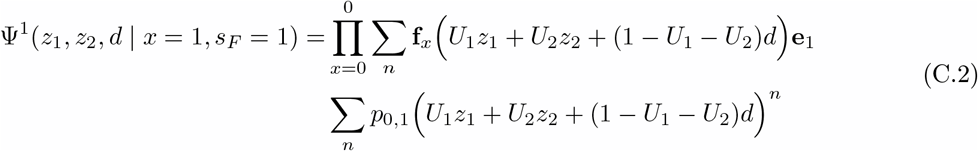

To find the joint probability of having *j*_1_ daughters in stage 1 and *j*_2_ in stage 2 and *m* losses, we take the multi-variate Taylor expansion:

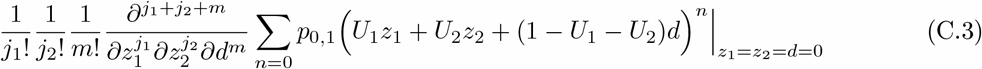

### Appendix C.2.1. Numerical examples

A couple of examples: Let *ω* = 3 age-classes 0,1,2; *k* = 2 stages 1,2; **U**, **U** = **I** _2_ and 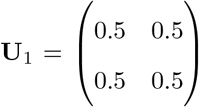 (assume no death). Suppose all newborns inherit stage of parent; Poisson reproduction only in age-classes 0 and 1: 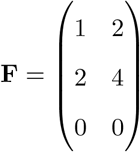 gives rates at which individuals of age (row) and stage (column) reproduce. The reproduction PGFs, *f*_*t,s*_(**z**), become e.g., 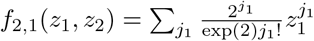 and 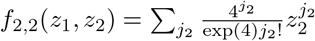.

#### Box 4

PGF for Daughters

Suppose that Focal is currently in age-class 2. Her offspring are represented by the Lifetime reproduction PGF:

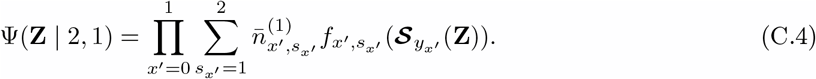

When Focal was in age-class 0, she is in stage 1. Her offspring must advance 2 time-steps: their PGF given by 𝒮_2_(**Z** | 1) = 0.5*z*_2,1_ +0.5*z*_2,2_ and 𝒮_2_(**Z** | 2) = 0.5*z*_2,1_ +0.5*z*_2,2_ (note however Focal only reproduces stage 1 offspring). The contribution towards Focal’s offspring PGF is:

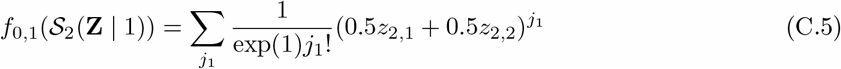

When Focal was in age-class 1 (*t* = 1), with probability [**U**_1_**U**_2_**e**_1_]_*j*_ = 0.5 she is in either stage *j* = 1, 2.

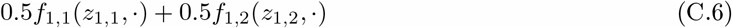

where 𝒮_1_(**Z** | *i*) = *z*_1,*i*_ because offspring inherit parent’s stage. Thus Focal’s contribution at age-class 1 is:

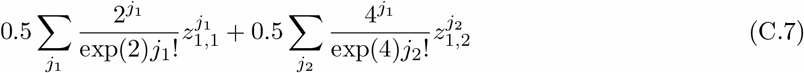

The PGF representing the age and stage of Focal’s duaghters becomes:

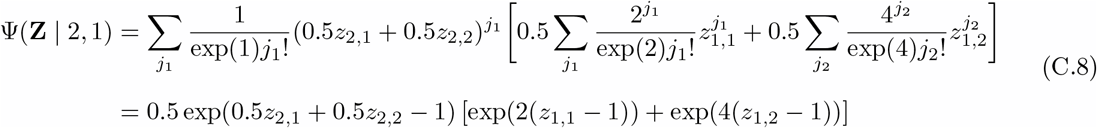

We find the joint probability, e.g., that Focal has *N*_2,2_ = 1 daughters of age 2 and stage 2, and *N*_1,1_ = 1 daughters of age 1 and stage 1 using the Taylor expansion:

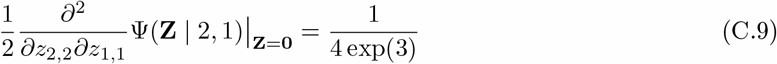

#### Box 5

PGF for granddaughters

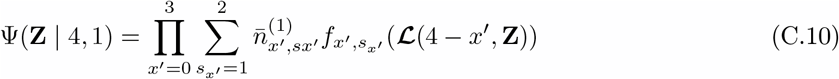

Focal’s stage-transitions are:

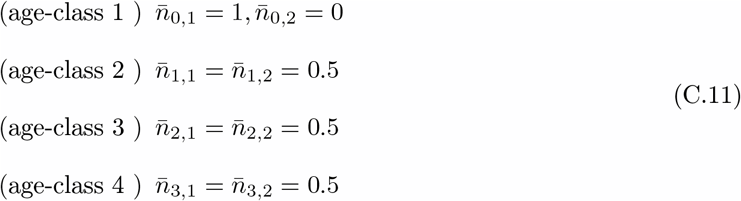

Daughters can only reproduce in first two age-classes. This implies grand-daughters can either be age-class 3 or 4. The granddaughters survival PGFs are:

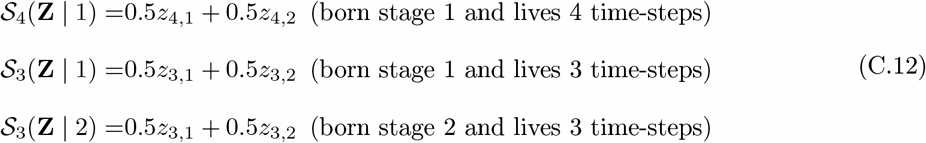

The lifetime reproduction PGFs of these daughters is, e.g., if daughter born into stage 1 is:

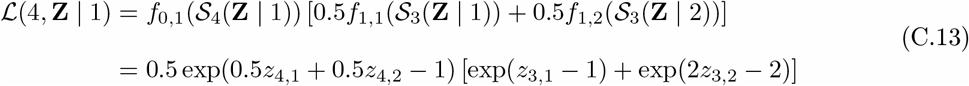

and similarly

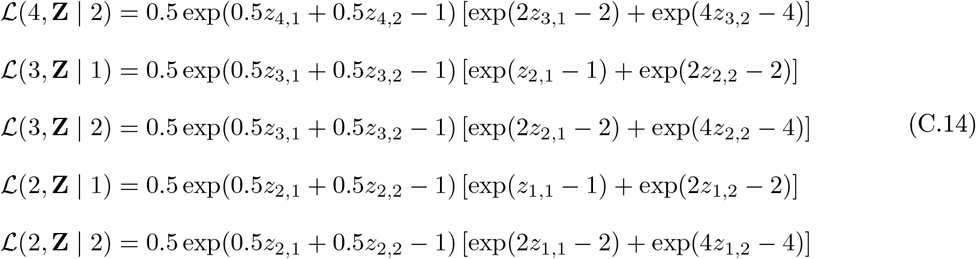

By age-class 4,

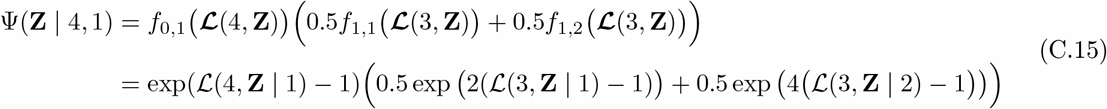

The joint probability that Focal has 1 grand-daughter of age-class 3 and stage 2, and one grand-daughter of age-class 2 and stage 1, is given by

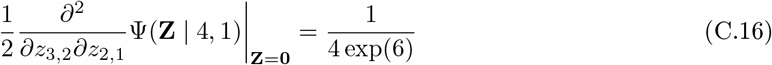

## Appendix D.

**Reducible projection matrices**

If the projection matrix is not irreducible, as in the parity-structured population example in text, there are a few ways in which we can infer Focal’s ancestry. In parity-structured populations, the stage variable records the number of children an individual has had. Transitions can only increase, never decrease. The projection matrix **Ã** therefore has a block lower-triangular structure. In this case, the Perron-Frobenius theorem does not guarantee a strictly positive eigenvector **w**; higher-parity states may have zero stable-population (and construction of the genealogical Markov chain involves division by *w*_*k*_ – undefined when *w*_*k*_ = 0).

### Appendix D.1. Restrict to reproductive age×stage combinations

One way around this problem is to restrict the genealogical decomposition to pathways that matter, i.e., the ones that can actually occur. Because the genealogical chain only needs to be defined on states actually visited by ancestral lineages, and since every ancestor must have been extant and reproductive, we can restrict attention to the set of reproductive states:

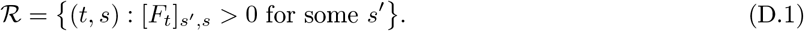

Let Ã| _ℛ_ denote the sub-matrix of Ã restricted to ℛ, for which Perron-Frobenious theory applies. Then the genealogical chain is then defined on ℛ. Under the assumption that Ã| _ℛ_ is irreducible with dominant eigenvalue *λ*| _ℛ_ and right Perron eigenvector | _ℛ_, the transition probabilities (updated from Eq (8)):

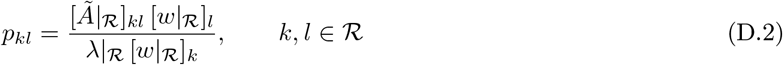

defines a valid Markov chain on ℛ leading to well-defined *π*(·) as per Eq (13). Note that this is our chosen method to define ancestral reproduction in the example in Section 4 of a population structured in parity.

### Appendix D.2. Numerical perturbation

A simpler, more pragmatic approach, but one that is less mathematically pleasing, is to allow non-zero (but virtually impossible) transitions between stages which ensure that the directed graph associated with the transition matrix is connected. This guarantees that the projection matrix is irreducible, that Perron-Frobenius holds, thereby meaning **w** is strictly positive with well-defined *p*_*k,l*_ for all *k, l*. Let **1** be of dimension *n* = *k*(*ω* +1): the total number of states. For small *ϵ >* 0, we apply a perturbation to the stage-transition matrix **Ũ** = [Ũ_*i,j*_] (see Eq (1)) to ensure it’s irreducible:

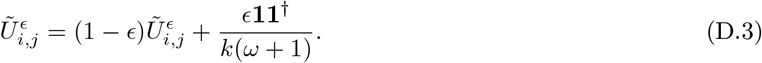

In the limit of *ϵ <<* 1, using the perturbed transition matrix 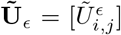 and 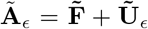 we construct the Markov chain transition matrix:

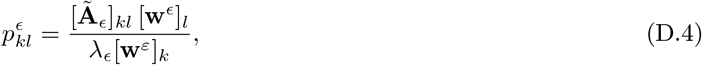

which is well-defined. Moreover, in the limit *ϵ* → 0 we recover well-defined *π*(·) as per Eq (13):

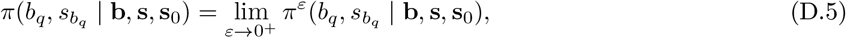

where *π*^*ε*^(·) denotes the ancestral joint probabilities computed from the perturbed Markov chain Eq (D.4). Furthermore, we see that if the limit in Eq (D.5) exists, then *π*(·) are supported on ℛ and coincide with the probabilities obtained from method Appendix D.1. That is, since 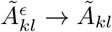 as *ϵ* → 0, and since 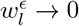 for any state *l* ∉ ℛ, the transition probabilities 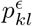 vanish for *l* ∈*/* ℛ. On ℛ, the restricted eigenvector **w**^*ϵ*^| _ℛ_ converges to **w**| _ℛ_ meaning 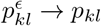.

## Appendix E

**Extending the framework to calculate living and dead kin by generation**

Here, we extend the PGF framework outlined in Section 6 to calculate the joint probabilities of having so many kin structured by generation which are living or dead. For simplicity, we marginalise over all ages and stages. Recall that in the [*g, q*]-kin convention, we the variable *g* represents the number of generations of descent from Focal’s *q*-th ancestor. As such, to count generation specific living kin of generation *i* = 0, …, *g* introduce the vector of dummy variables **z**_*g*_ = (*z*_1_, …, *z*_*g*_), i.e., *z*_*i*_ counts the number of living kin in generation *i* (over all ages and stages). We introduce the vector of dummy variables **d**_*g*_ = (*d*_1_, …, *d*_*g*_) where *d*_*i*_ counts the numbers of dead kin of generation *i*. The recursive composition of lifetime reproductive PGFs which count deaths in Eq (22) is adjusted so that ℒ^(*i*)^ now includes the vectors of dummy variables **z**_*g*_ = (*z*_1_, …, *z*_*g*_) and **d**_*g*_ = (*d*_1_, …, *d*_*g*_), where *z*_*i*_ and *d*_*i*_ respectively count the living and dead kin of generation *i* ∈ {1, …, *g*}. We define ℒ^(*g*)^ as the PGF for a lineage starting from a newborn of generation *g*, which tracks the survival and reproduction of that individual and all their subsequent descendants up to the terminal generation *g*.

The PGF for an kin of generation *g* of descent tracks only their own survival or death at age *h*; they do not produce further kin (in our [*g, q*] convention):

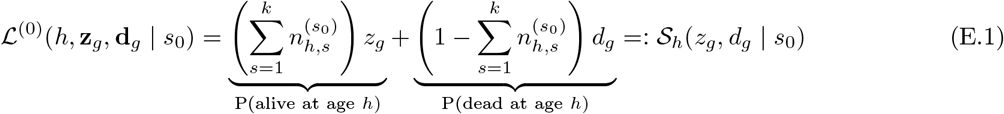

While (0 *< i < g*) the PGF tracks the individual’s survival (*z*_*i*_), their death (*d*_*i*_), and production of offspring who start a new lineage of generation *i* + 1:

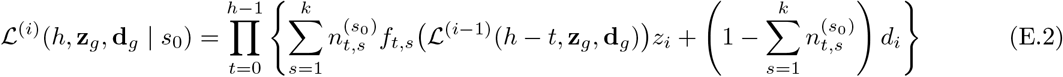

For example, consider Focal’s descendants. When Focal’s daughter is alive she produces granddaughters (using ℒ^(0)^), while her own survival is tracked by *z*_1_. When Focal’s daughter dies, her survival is tracked by *d*_1_, and her production stops.

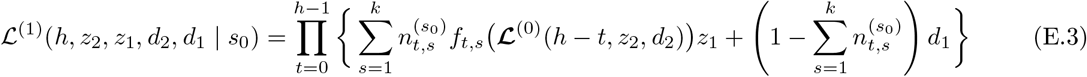

Then, substituting Eq (E.3) into Eq (23) recovers a PGF for the number of granddaughters (*g* = 2) when Focal is *x*+1, denoted Ψ^*g*=2^(**z**, *z*_2_, *z*_1_, *d*_2_, *d*_2_ | *x*+1, *s*_*F*_), which also keeps track the number of Focal’s daughters (*g* = 1) which are dead.

In text we consider an example where Focal experiences *j* living granddaughters (*g* = 2 descendants) but whose mothers (the *g* = 1 descendants) have died. Consider such *orphaned* granddaughters (recall we assume a one-sex population). By setting *z*_1_ = 0 we consider the case of no living daughters, and setting 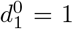 we marginalise (count over) over all dead daughters. By setting 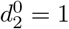 we marginalise over dead all granddaughters. In this case, the PGF which calculates the probable numbers of granddaughters in the case where all of Focal’s daughters are dead is:

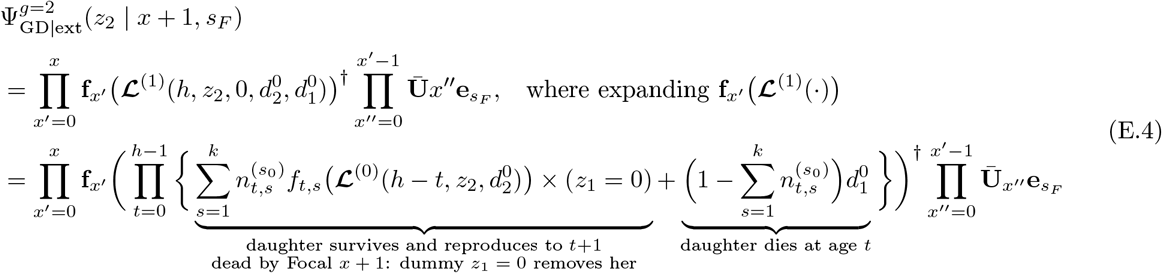

where *h* = *x* + 1 − *x*′ is the age the daughter would have reached at Focal’s present. The first term inside the inner product (the summation) represents the daughter surviving. Because we are looking for the case of total daughter extinction, any instance whereby daughter is still alive at any age *t < h* is multiplied by *z*_1_ = 0, effectively removing those branches. The dummy variable *d*_1_ represents the daughter’s death: 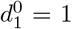 marginalises over the death events. The granddaughters’ survival (term 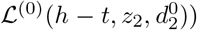, nested inside the daughter’s fertility PGF *f*_*t,s*_, counts the granddaughters produced by the daughter before her death, and tracks their survival to the present.

Alternatively, we treat Focal’s granddaughters who have lost a mother individually. That is, we find the probable numbers of granddaughters who are alive but whose own mothers (Focal’s daughters) are dead (but not necessarily all of Focal’s daughters are dead). Let 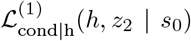 represent the PGF of the complete reproduction of Focal’s daughter *who survives to age h*:

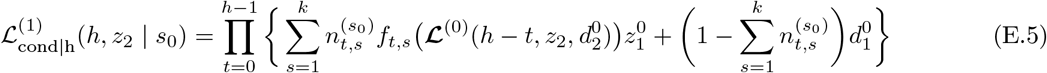

where 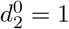 since we count living granddaughters, 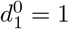 as we condition on daughter alive at age *h*. Orphan granddaughters are counted when the daughter’s are dead by the end of the interval *h*, in which case daughter contributes her accumulated granddaughters. Let 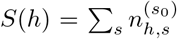 be the probability the daughter is alive at when at age *h*. The PGF for one daughter’s orphaned offspring is then 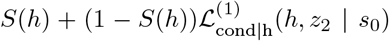. Substituting this into the PGF for Focal’s total descendants, we see for Focal at age *x* + 1, the PGF for the number of granddaughters whose own mothers are deceased:

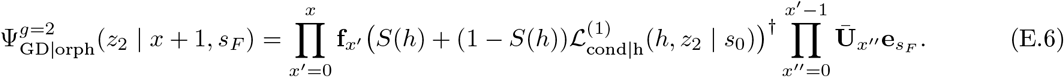

Fig 8 of Section 4.4 in text plots the probabilities that Focal has exactly *j* orphaned granddaughters, found by the coefficients of 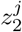:

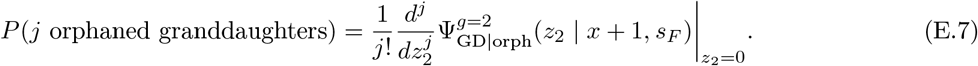

## Appendix F.

**Extracting numerically**

For a scalar PGF Ψ(*z*) =∑ _*n*_ *p*_*n*_*z*^*n*^, the probabilities *p*_*k*_ are found through the Cauchy integral:

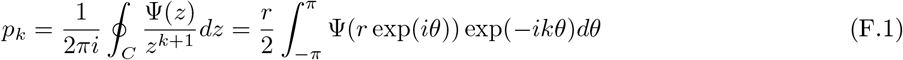

where *r* is the radius of the closed curve *C*. Since by definition *z* ≤ 1, integrating over the unit circle suffices to enclose all poles of Ψ(*z*). As such:

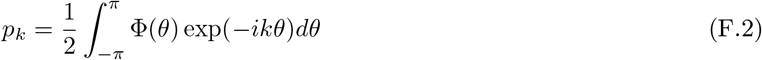

where Φ(*q*) = Ψ(exp(*iq*)) is characteristic function, is a Fourier expansion. Numerical integration allows us to approximate:

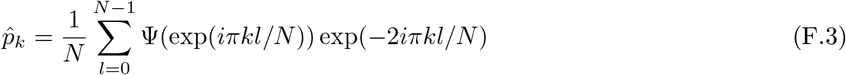

where *N* is transform length, or the number of distinct points. The above is simply a Discrete/Fast Fourier transform. This approach is taken in our computational implementation.

The multivariate PGF is defined over the matrix of dummy variables **Z**:

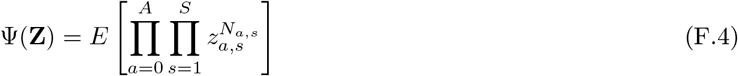

and the joint probabilities of state-specific numbers **n** = {*n*_0,1_, …, *n*_*A,S*_} is found by:

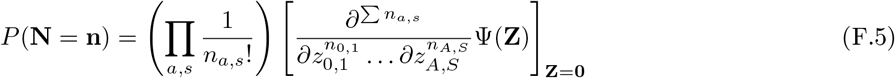

mapping the PGF to the probability mass function (PMF) across the age-stage range. Numerically, employing FFTs, let the state coordinate be

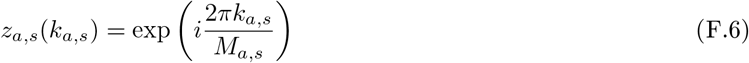

where *M*_*a,s*_ is the transform length (maximum count + 1) for that specific state. Then

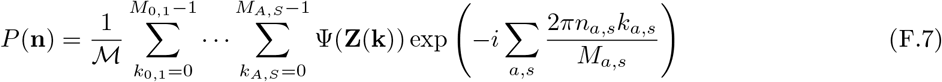

where ℳ = Π *M*_*a,s*_ is the total number of grid points.

## Appendix G

**Reproducing Caswell [18]**

Here we compare the expected numbers of kin by stage obtained from the above framework, to the already published parity-model of Caswell [18]. Regarding daughters, the PGF model recovers precisely same expected numbers by parity level, as shown in Fig G.1:

**Figure G.1:**
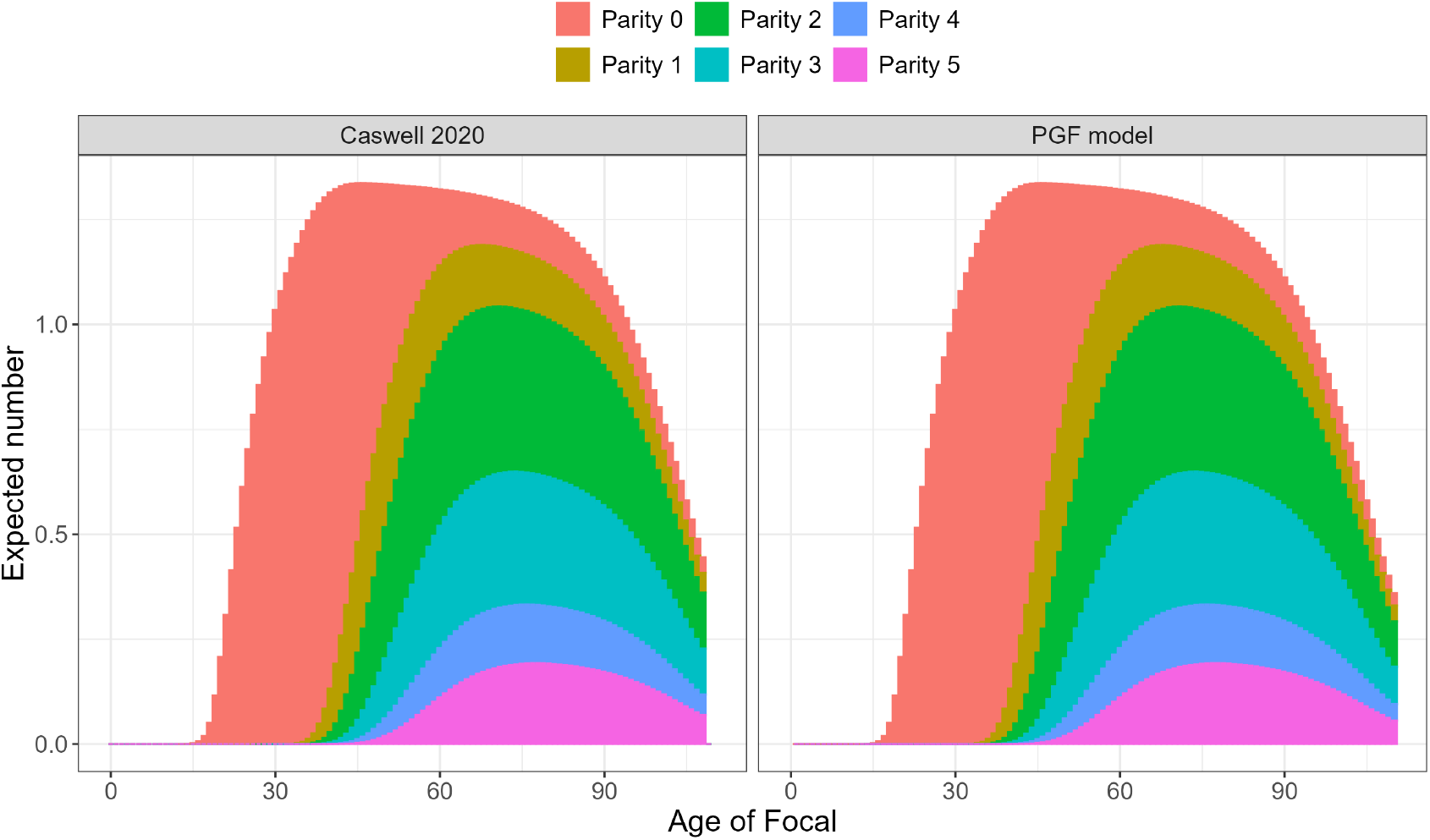
Comparing the expected numbers of daughters, by parity level of daughter, and by age of Focal. Author’s own implementation of the model of Caswell [18] using data therein.

Regarding sisters, the PGF model recovers a similar qualitative pattern, but overall lower levels, for the expected numbers by parity level; Fig G.2.

**Figure G.2:**
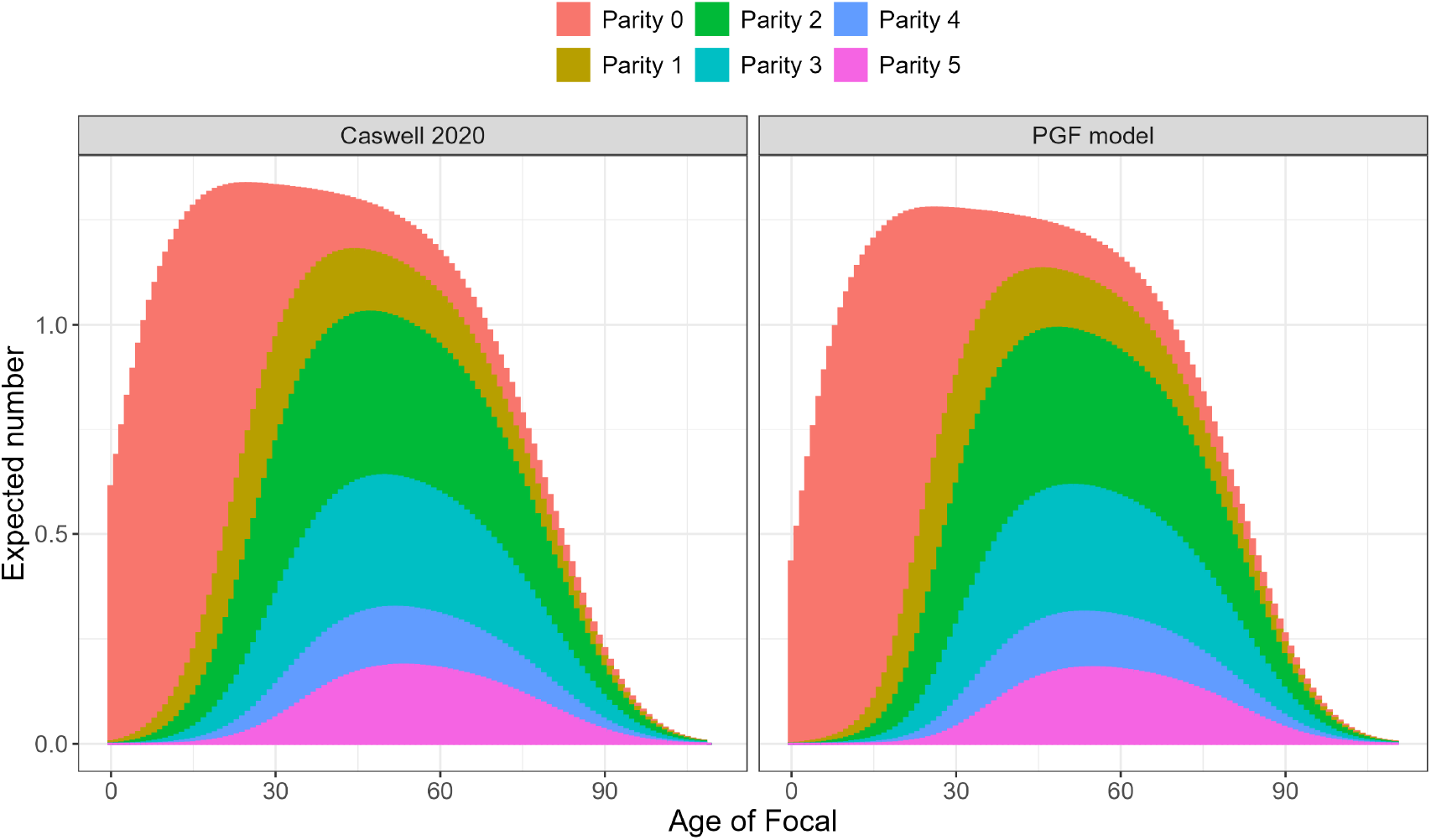
Comparing the expected numbers of sisters, by parity level of sister, and by age of Focal. Author’s own implementation of the model of Caswell [18] using data therein.

## Footnotes

1 We note a restriction in the model here and address possible solutions in Appendix D

